# Dormant phages communicate to control exit from lysogeny

**DOI:** 10.1101/2021.09.20.460909

**Authors:** Nitzan Aframian, Shira Omer Bendori, Stav Hen, Polina Guler, Avigail Stokar-Avihail, Erica Manor, Kholod Msaeed, Valeria Lipsman, Ilana Grinberg, Alaa Mahagna, Avigdor Eldar

**Affiliations:** Shmunis School of Biomedicine and Cancer Research, Faculty of Life Sciences, Tel-Aviv University, Tel-Aviv, 69978, Israel; Dept. of Molecular Genetics, Weizmann Institute of Science, Rehovot, 7610001, Israel; Dept. of Plant and Environmental Sciences, Weizmann Institute of Science, Rehovot, 7610001, Israel

## Abstract

Temperate bacterial viruses (phages) can transition between lysis - replicating and killing the host, and lysogeny - existing as dormant prophages while keeping the host viable. It was recently shown that upon invading a naïve cell, some phages communicate using a peptide signal, termed arbitrium, to control the decision of entering lysogeny. Whether communication can also serve to regulate exit from lysogeny (known as phage induction) remains unclear. Here we show that arbitrium-coding prophages continue to communicate from the lysogenic state by secreting and sensing the arbitrium signal. Signaling represses DNA-damage dependent phage induction, enabling prophages to reduce induction rate when surrounded by other lysogens. We show that the mechanism by which DNA damage and communication are integrated differs between distantly related arbitrium-coding phages. Additionally, signaling by prophages tilts the decision of nearby infecting phages towards lysogeny. Altogether, we find that phages use small molecule communication throughout their entire life-cycle to measure the abundance of lysogens in the population, thus avoiding wasteful attempts at secondary infections when they are unlikely to succeed.

## Introduction

Temperate phages are able to alternate between two different states - a lytic state where they replicate, kill the host and release their progeny into the environment, and a dormant, lysogenic state, during which they typically integrate into the bacterial chromosome as prophages and replicate along with it. Temperate phages are thus faced with a critical decision between lysis and lysogeny at two points within their full life-cycle - a decision of whether to enter lysis or lysogeny upon infection, and a decision made from the lysogenic state whether to persist as prophages or exit lysogeny through phage induction. In the highly studied phage λ, the lysis-lysogeny decision upon infection is controlled by phage density through detection of co-infection ^1,2^, while DNA damage to the host cell serves as a trigger for induction ^3^. Other factors such as the physiological state of the cell were also shown to influence the development of phage λ ^4^.

Recently, several groups of phages infecting the genus *Bacillus* were shown to employ an alternative mechanism for regulating the lysis-lysogeny decision upon infection, which is based on small-molecule communication. These phages encode for the RRNPP type arbitrium communication system ^5–7^. Upon entry into the cell, the phage-encoded intracellular receptor AimR and the pre-AimP arbitrium pre-peptide are expressed. Pre-AimP is cleaved upon secretion to form the mature arbitrium peptide, AimP, which can be transported back into the cell by the general oligopeptide permease (Opp) system. In the absence of mature AimP, AimR acts as a transcription factor to activate the transcription of the *aimX* regulatory RNA., Intracellular mature AimP interacts with AimR and prevents it from activating the expression of *aimX* ^8–11^. In SPβ-like phages (clade II arbitrium systems), *aimX* most likely promotes the lytic state through uncharacterized trans-interactions. In contrast, in many other arbitrium clades, *aimX* is transcribed in an antisense direction to a putative phage repressor and seems to operate in *cis* to reduce its transcript level ^6^.

In total, the arbitrium system biases infecting phages towards lysogeny at late rounds of infection, when the arbitrium peptide reaches high concentrations and phage density is high. In this sense, small-molecule communication seem to provide a similar phage density-dependent effect on lysis-lysogeny decision-making to that provided by the coinfection-sensing mechanism employed by phage λ^12,13^. It is therefore unclear whether there are critical population-level differences between small-molecule signaling and coinfection sensing. One attractive hypothesis we consider here, is that small-molecule signaling can continue from the lysogenic state, allowing prophages to inform other phages of their presence, a function which could not be achieved by coinfection sensing^14^. The frequency of lysogens in the population is valuable information from the phage perspective – lysogens are often immune to secondary infections, a phenomenon termed superinfection exclusion ^15^. Therefore, the probability of failed infection attempts increases with lysogen frequency and the adaptive value of lysis decreases accordingly.

Here we show that prophages possessing the arbitrium system indeed continue to communicate intercellularly from the lysogenic state. We demonstrate that signaling inhibits the process of phage induction triggered by DNA damaging agents, and that signaling by prophages influences infection dynamics. Overall, we show that signaling from the lysogenic state allows phages to avoid lysis when surrounded by lysogens.

## Results

### Prophages communicate by secreting and sensing AimP

In contrast to co-infection based mechanisms of sensing phage density, the arbitrium system may enable phages to communicate their presence from the dormant prophage (lysogenic) phase ^14^. The activity of the arbitrium system during lysogeny and its possible impact have not been previously characterized, although global transcriptome analysis of the *B. subtilis* lab strain 168, which hosts the arbitrium-carrying SPβ prophage, indicates that *aimR, aimP* and *aimX* are transcriptionally active ^16,17^.

To investigate the possibility of communication during lysogeny, we first asked whether prophages can sense the arbitrium peptide during the lysogenic phase. To this aim, we introduced into an ectopic chromosomal locus an *aimX*-YFP transcriptional reporter containing the *aimX* promoter and part of the *aimX* transcript in an operon with three consecutive YFP genes (methods). Comparison to reporter fluorescence in non-lysogens indicated that *aimX* is expressed in lysogens during growth in LB medium (Fig. 1A). Expression was reduced upon addition of the mature SPβ AimP arbitrium peptide (GMPRGA,SPβ AimP) but not upon addition of the orthogonal mature AimP of phage ϕ3T (SAIRGA, ϕ3T AimP) ^5^, suggesting that AimR continually activates *aimX* during lysogeny and responds to AimP. Indeed, deletion of *aimR* from the prophage completely abolished *aimX* expression.

**Figure 1:**
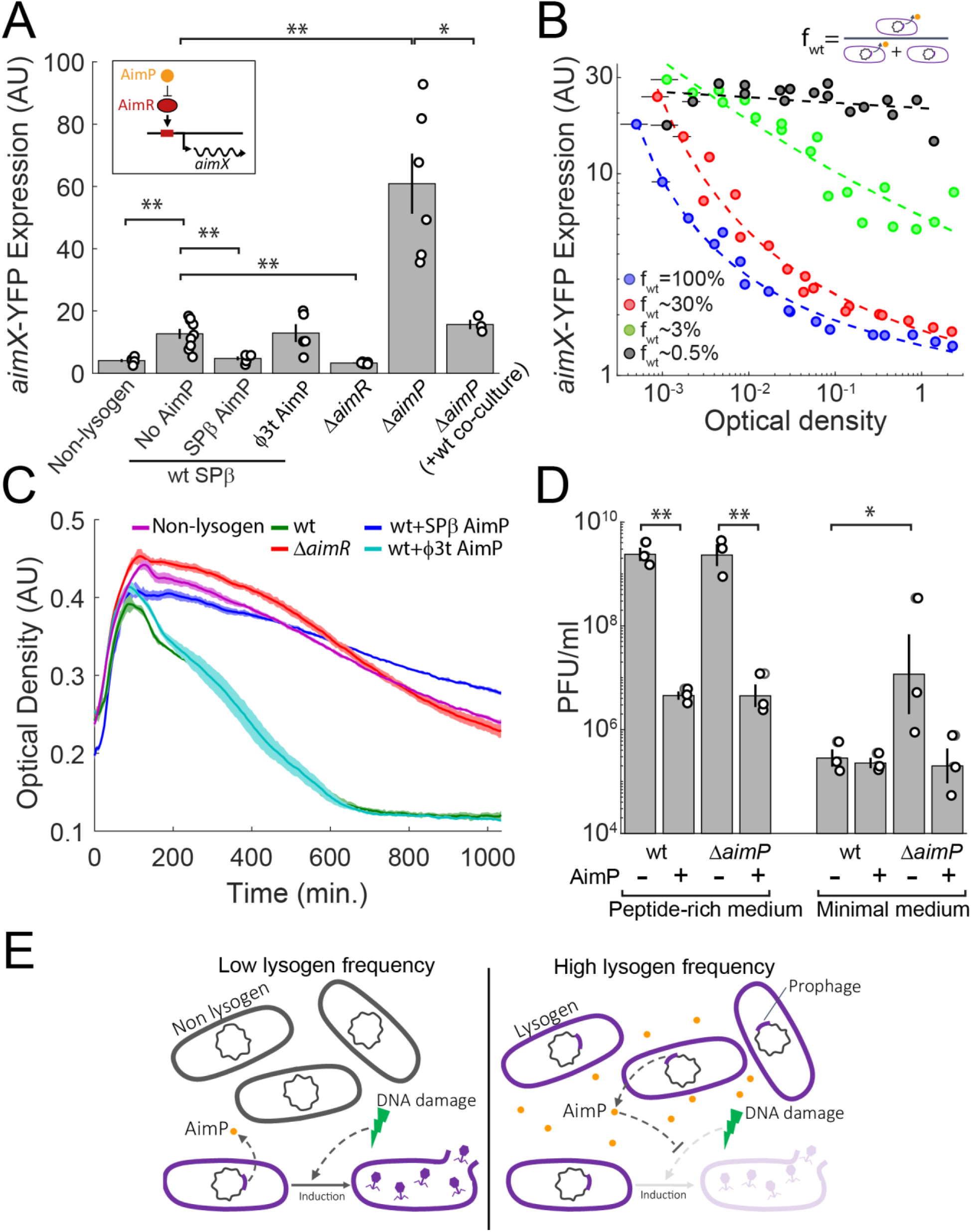
Arbitrium signaling is active during lysogeny and represses DNA damage-dependent prophage induction. (A) Mean expression of an *aimX*-YFP reporter in LB at OD_600_ = 0.3 for the various genotypes and conditions described. Empty circles represent median expression level of individual biological repeats. Error bars represent standard errors. The rightmost bar is the expression of a Δ*aimP* mutant lysogen in 1:1 coculture with wild-type lysogen (Δ*aimP* mutant strain was marked by BFP to distinguish it from the wild-type during flow-cytometry). * 0.005 < *p* < 0.05, \*\**p* < 0.005. (n=5,6 repeats of each condition, paired t-test). Each individual measurement is the median of YFP obtained by flow-cytometry over >10^4^ cells. Measurements were taken on different days. AimP signal was added at a concentration of 10μM. (B) Expression of a wild-type lysogen carrying an *aimX*-YFP reporter in co-culture with a *ΔaimP* strain as a function of optical density and frequency. Different colors show different approximate frequencies (f) of the wild-type as indicated in the legend. Dashed lines show the best fit of data for each frequency used to illustrate the trend (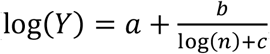, where *n* is the cell density). Low optical densities (<0.02) are calculated by back-extrapolation based on time of measurement and measured growth rate in minimal medium (50±8.5 minutes, Supplementary Fig. 2). Error in calculation of apparent optical density are marked by horizontal line of the appropriate color whenever this error is larger than the marker (very low optical densities). (C) Mean optical density (dark color) and its error (lighter shade) as a function of time (minutes) for the relevant strains and condition (see legend, Table S1), taken using a plate-reader. 0.5 μg/ml Mitomycin C (MMC) was added at time 0 at an OD_600_=0.3. Results are average of three technical repeats. See additional biological repeats and statistical analysis in Supplementary Fig. 1. (D) Plaque forming units (PFU) for cultures induced with MMC at OD_600_=0.3 and grown in peptide-rich medium (LB, lysate taken 120 minutes after addition of MMC) or in minimal medium (SMM, lysate taken 150 minutes after addition of MMC). Empty circles represent separate biological repeats and error bars represent standard error of the log. * 0.005 < *p* < 0.05, \*\**p* < 0.005. (n=3, paired t-test). (E) an illustration of the basic findings of this section – at low lysogen frequency DNA damage leads to prophage induction. At high lysogen frequencies, the lysogen-secreted arbitrium signal inhibits induction. Lysogens are marked by purple color indication of their outline and of the prophage within their chromosome.

In order to test whether the arbitrium peptide is not only sensed, but also produced by lysogens, we measured *aimX* expression in lysogens deleted in *aimP* and grown in LB (Fig. 1A). Deletion of *aimP* led to a substantial increase in *aimX*-YFP expression, implying that AimP is continuously produced during lysogeny and inhibits activation by AimR. To show that AimP is not only produced but also secreted, we co-cultured differently marked wild-type and Δ*aimP* lysogens (Fig. 1A, B). YFP Expression levels of the *aimX*-YFP reporter in the Δ*aimP* lysogens were reduced to levels of the wild-type, as expected from a secreted peptide. Overall, these results indicate that prophages communicate out of lysogeny by a process of quorum sensing – the arbitrium peptide is secreted, and sensed by surrounding lysogens.

To further explore the behavior of the arbitrium system in lysogeny, we monitored the expression of an *aimX*-YFP transcriptional reporter at different densities during growth in minimal medium (Fig 1B). As expected, the *aimX* reporter was highly expressed at very low densities, but was gradually turned off starting at a cell density of 10^5^ cells/ml (equivalent to an effective optical density of ∼10^−3^, methods). This shows that the arbitrium system is highly sensitive compared to other RRNPP quorum-sensing systems, where an effect can be observed only at optical densities which are more than 10 times higher^18,19^. As a further indication for the sensitivity of the system, we found that at very low cell densities addition of 1-10pM of signal to a Δ*aimP* mutant was sufficient to partially repress the *aimX*-YFP transcriptional reporter (Supplementary Fig. 2). This stands in contrast to reported levels of 1-10nM in other RRNPP systems ^20^. Notably, much higher concentrations are needed at high cell densities, presumably since cells uptake and degrade the peptide signal (Supplementary Fig. 2) ^21,22^.

### Communication controls prophage induction

Next, we set out to explore the phenotypic response controlled by prophage communication. DNA damage triggers induction in many distantly related phages, including SPβ-like phages ^23,24^. This strategy may be adaptive to the phage as DNA damage strongly reduces host viability and increases the relative benefit of horizontal transfer ^25^. Since communication between infecting phages controls the decision to enter lysogeny, we hypothesized that communication between dormant prophages may control the decision to exit lysogeny. DNA-damage dependent induction may become maladaptive when a lysogen is surrounded by other lysogens. This is especially true when free phages are able to adsorb to lysogens but cannot complete productive infections. Our results suggest that SPβ lysogens can indeed act as sinks of SPβ phages (Supplementary Fig. 3).

In order to understand the interplay between the inputs of signaling and DNA damage in SPβ induction, we generated DNA damage by adding Mitomycin C (MMC) to a rich medium in which lysogens were grown. Phage induction was monitored both by tracking bacterial density (Fig. 1C, Supplementary Fig. 1) free phage production (Fig. 1D). Non-lysogens exposed to MMC showed arrested growth and a slow reduction in OD as expected in the face of DNA damage ^26^, while lysogens experienced a sharp population collapse due to phage induction (Fig. 1C, Supplementary Fig. 1). We note that *B. subtilis* lab strains contain an additional lytic element, the degenerate phage-derived bacteriocin, PBSX ^27^. Our measurements were therefore done in a genetic background where PBSX is inactivated by mutations (methods).

We found that addition of ectopic SPβ AimP along with MMC to SPβ lysogens reduced phage production and prevented population collapse (Fig. 1C, D), while addition of a peptide from a different arbitrium system, ϕ3T AimP, had no discernible effect (Fig. 1C). This indicates that arbitrium signaling dominantly represses the activation of induction in response to DNA damage. As expected from this regulation, addition of MMC to Δ*aimR* lysogens did not lead to massive cell lysis and population collapse (Fig. 1C).

Since we established that AimP inhibits prophage induction, we expected induction of *ΔaimP* prophages to be even more rapid compared to the wild-type. However, when grown in LB, induction rates did not differ between these two strains (Fig. 1D Supplementary Fig. 1). One possible explanation is that expression levels of *aimX* in WT lysogens grown in LB may not be sufficiently low to significantly prevent induction. We reasoned that this might be due to low intracellular concentrations of the arbitrium peptide resulting from competition over the Opp transporter with other peptides, which are abundant in LB medium at this growth stage ^21^. Supporting this hypothesis, we found that while *aimX* is expressed at mid-log phase in LB (Fig. 1A), expression in peptide-free minimal medium is completely shut down at this growth stage (Fig. 1B). Additional physiological factors may also contribute to the effect of the medium on phage decision-making, as has been described for other phages ^28^. In agreement with *aimX* expression, addition of MMC to Δ*aimP* lysogens grown in minimal medium led to significantly more induction compared to WT lysogens (Fig. 1D). Addition of ectopic AimP along with MMC to Δ*aimP* lysogens, reduced induction rates to WT levels. As expected given the complete suppression of *aimX* expression in minimal media (Fig. 1B), induction of WT prophages was not affected by supplementing minimal medium with ectopic AimP.

Since the frequency of lysogens in the population is a crucial variable for the probability of successful future infections, we expected expression patterns of *aimX* to depend on the frequency of signal producers. To test this, we co-cultured differentially marked wild-type and Δ*aimP* lysogens in minimal medium, and tracked *aimX*-YFP reporter expression in WT lysogens (Fig. 1B). We found that under these conditions, ∼3% of WT lysogens in the population were sufficient to partially suppress *aimX* expression by mid-log phase, while a fraction of ∼30% led to strong suppression. In contrast, when WT lysogens make up <1% of the population, high levels of *aimX* expression are maintained even at high densities. To conclude, we find that prophages communicate during lysogeny to inhibit induction when they sense they are surrounded by other lysogens, as indicated by high concentrations of the arbitrium peptide (Fig. 1E). This strategy may allow phages to avoid induction under conditions where their progenies are likely to encounter established lysogens rather than permissive bacteria.

### Communication inhibits induction through different mechanisms in different phages

To gain further insight into the inhibition of induction by arbitrium signaling, we used RT-PCR to measure the transcript levels of several SPβ phage genes which come into play at different stages of prophage induction (Fig. 2A). A rise in transcription levels of *yorB*, which is regulated directly by LexA, the bacterial SOS repressor ^23^, were observed just 20 minutes after MMC addition and were not strongly affected by the presence of the arbitrium signal. However, transcription levels of *sprB*, which controls prophage excision ^29,30^, rose only later (45 minutes after MMC addition) in the presence of DNA damage, but only when the medium was not supplemented with the arbitrium peptide. Transcription of the later expressed genes coding for phage RNA-polymerase (*yonO*), putative phage capsid (*yonH*) and phage lysin (*cwlP*) ^31,32^ were similarly activated by DNA damage and inhibited by addition of the signal. Overall, these results suggest that the arbitrium signal prevents the expression of the phage excision module without shutting down the general SOS response regulon. This indicates that when sensing lysogens, prophages remain dormant even in the occurrence of DNA-damage, which allows their bacterial host to attempt repair.

**Figure 2:**
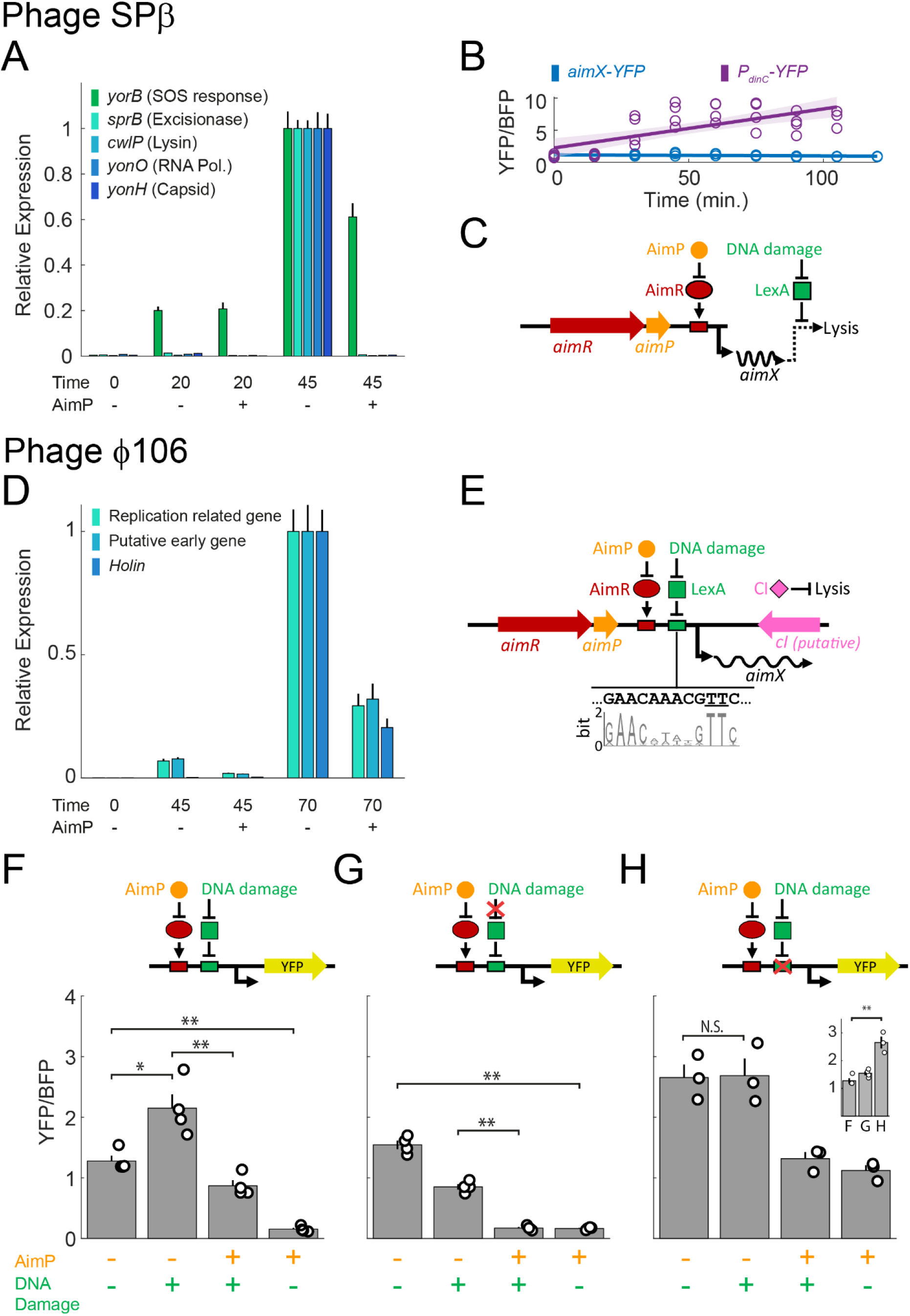
Arbitrium signal and DNA damage are differentially integrated in different phages. Shown are results for phage SPβ (A-C) and ϕ106 (D-H). (A) Relative phage transcript expression of several SPβ genes (legend) measured using RT-PCR for different times after addition of 0.5μg/ml of MMC in the presence or absence of 10μM of ectopic SPβ AimP. Data is normalized to 45 minutes after MMC addition in the absence of peptide. Shown are averages of three biological replicates, each with two technical replicates. Error bars represent 95% confidence interval (methods). (B) Expression of the *aimX*-YFP reporter of phage SPβ (blue, slope estimate *p* = 0.08, *n* = 26) as a function of time after addition of MMC, taken using flow cytometry. For each measurement, median YFP levels are divided by median BFP levels from a constitutive P_veg_- BFP reporter (methods). Also shown are the YFP expression of transcriptional reporter of a known LexA-repressed gene, P_*dinC*_-YFP divided by the same BFP reporter (purple, slope estimate *p* = 2 × 10^‒5^, *n* = 29). Solid lines and lighter shade represent linear fit of the corresponding data and its error. (C) Illustration: In phage SPβ, LexA-mediated regulation of phage induction occurs downstream of *aimX* expression. (D) As in (A), but for three ϕ106 associated genes (legend, methods). Data is normalized to 70 minutes after MMC addition in the absence of peptide. (E) Analysis of the ϕ106 arbitrium module identified a LexA binding site within the *aimX* locus and suggests a direct regulation of *aimX* expression by LexA and putative indirect regulation of repressor activity through the previously suggested sense-antisense mechanism^6^. Shown below the illustration are the identified LexA binding site and below it the sequence logo of LexA binding motif. Underlined are two highly conserved thymines which are replaced by adenines in the defective LexA binding mutant. (F-H) Verification of LexA effect on *aimX*. Shown are expression of an *aimRP*X _*ϕ106*_-YFP reporter of phage ϕ106 arbitrium module, divided by a constitutive BFP reporter. Data is taken 90 minutes after the conditional addition of ϕ106 AimP and MMC (see Supplementary Fig. 7 for temporal data). Three different strains were assayed: (F) A strain carrying an unmodified arbitrium module. (G) A strain carrying an unmodified arbitrium module expressing a high level (Supplementary Fig. 8) of a non-cleavable dominant negative variant of LexA ^23^. (H) A strain carrying a modified arbitrium module mutated in two critical base-pairs of the LexA binding motif (shown in methods). Inset: comparison of the first column (no MMC or AimP) in panels F, G, H. Illustrations above each graph show the proposed regulation. Red X point to the changes in regulation obtained by the modifications in (G, H). At least three measurements (open circles) were taken at different days. Shown are means and standard errors for each strain and condition. * 0.005 < *p* < 0.05, \*\**p* < 0.005. N.S – non-significant (n=3,4, paired t-test).

DNA damage and arbitrium signals could be integrated at multiple different stages to yield the final decision of induction. To examine whether DNA damage acts upstream of the arbitrium communication system and affects its expression, we monitored *aimX* expression levels through our *aimX*-YFP fluorescent reporter. As cells do not divide under DNA damage, their total fluorescence may rise. We controlled for this effect by dividing the YFP by the BFP expression of a constitutive BFP reporter (methods, Supplementary Table 1). We found this relative *aimX*-YFP expression to be independent of DNA damage (Fig. 2B, Supplementary Fig. 4). The DNA damage induced gene *dinC* ^23^ was used as a positive control, which indeed showed an increase in expression in response to DNA damage. *aimX*-YFP expression was generally higher in a Δ*aimP* background, and in fact showed a slight reduction in the relative expression, opposite to an expected direct effect of DNA damage on *aimX* (Supplementary Fig. 4). The lack of significant effect on *aimX*-YFP was further verified by measuring the transcript level of our *aimX*-YFP reporter using RT-PCR (*aimX* on its own is too short for accurate quantification using standard RT-PCR methods, Supplementary Fig. 5). Altogether, these results indicate that while *aimX* transcription is regulated by AimP, as shown above, it does not respond to DNA damage. These results are further corroborated by publicly available expression data which monitored gene expression upon addition of MMC in the *B. subtilis* 168 lab strain (Supplementary Fig. 6) ^17^. Therefore, integration of DNA damage and communication in phage SPβ occurs downstream of *aimX* expression (Fig. 2C).

We next wanted to explore the generality of our findings in other arbitrium coding phages. Arbitrium systems are found in other phages of the genus *Bacillus*, and phylogenetic characterization identified 10 clades of AimR, with SPβ-like phages encoding clade 2 receptors ^6^. To investigate whether other phages besides SPβ, also regulate phage induction using arbitrium communication, we initially examined the SPβ-like phage ϕ3T, which also harbors a clade 2 arbitrium system. As in SPβ lysogens, DNA-damage caused by addition of MMC lead to population collapse of ϕ3T lysogens, which was inhibited specifically when ectopic ϕ3T arbitrium peptide (SAIRGA, ϕ3T AimP) was added to the medium (Supplementary Fig. 9).

As explained above, in some arbitrium clades, *aimX* overlaps a putative phage repressor and is expressed in the opposite direction, suggesting that it inhibits repressor expression through *cis*-antisense regulation^6^. We wondered whether signaling prevents phage induction in such an architecture as well. *B. subtilis subsp. inaquosorum* strain KCTC13429 contains a phage similar to phage ϕ105 in its lytic genes but altered in its lysogenic domain (Supplementary Fig. 10). This phage family, which we termed ϕ106 codes for an arbitrium system with clade 1 receptor, and has the abovementioned architecture of sense-antisense arrangement between the arbitrium system and a putative repressor (Fig. 2E, Supplementary Fig. 11). We used RT-PCR to monitor the induction of this phage upon addition of MMC, either with or without the putative arbitrium AimP signal of this system (DPPVGM). We monitored three genes; two putative early genes - a transcription factor and a replication organizer homolog, and a late-stage holin homolog. We found that addition of MMC strongly activated these genes. Addition of the AimP signal together with MMC reduced the expression of these genes by a factor of five compared to addition of MMC alone (Fig. 2D). This is a substantial level of repression, though weaker than that observed for SPβ (Fig. 2A). These results indicate that repression of induction by signaling may be a general theme in arbitrium systems.

The ϕ106 phage of *B. subtilis subsp. inaquosorum* does not infect the *B. subtilis* 168 lab strain or any of its derivatives, preventing us from studying this phage as a whole in this model strain. To further understand the mechanism by which DNA damage and arbitrium signals are integrated in this phage, we cloned the ϕ106 arbitrium system together with part of *aimX* fused to a YFP gene (*aimRPX*_*ϕ106*_-YFP, Supplementary Fig. 7) into a background strain which lacks all other inducible elements and contains a constitutive BFP reporter (Supplementary Fig. 7). We then examined *aimX* expression by monitoring the mean YFP/BFP ratio in the population under different environmental perturbations (Fig. 2F). We found that the addition of MMC increased the relative *aimX* reporter expression. Addition of peptide decreased both the basal expression without MMC and the MMC-induced expression. These results therefore indicate that in this system, in contrast to phage SPβ, DNA damage is integrated with signaling upstream of *aimX* expression (Fig. 2E, top).

Upon inspection of the *aimX* sequence, we noted the existence of a nucleotide sequence with high similarity to the consensus binding site of the DNA damage-responsive SOS repressor, LexA (Fig. 2E) ^33^. This led us to the hypothesis that DNA damage regulation in this phage occurs at the level of *aimX* transcription. To validate the role of LexA and the putative LexA binding site in the transcriptional regulation of *aimX*, we constructed two perturbed strains. In the first, we overexpressed a dominant negative non-cleavable allele of LexA, known as LexA(ind-) (methods)^24^. This allele is expected to prevent the expression of LexA-dependent genes under DNA damage conditions (Fig. 2G, top). This was verified by examining its effect on a *dinC* reporter (Supplementary Fig. 8). Monitoring *aimRPX*_ϕ106_-YFP expression in a background of this dominant negative allele, we found that addition of MMC to the cells did not increased *aimX* expression (Fig. 2G). In-fact we observed a slight decrease in reporter activity, most likely due to changes in cellular response to DNA damage in the LexA dominant allele background, as reflected in the changes to constitutive gene expression in this background (Supplementary Fig. 8). Addition of peptide still led to repression of *aimX* expression either with or without MMC. These results suggest that LexA mediates the DNA damage regulation of *aimX* expression.

Second, to validate the impact of the putative LexA binding site, we mutagenized the *aimX*-YFP reporter by modifying two critical bases in the binding site (Fig. 2E). This mutation is expected to derepress *aimX* expression irrespective of DNA damage (Fig. 2H, top). Indeed, we found that gene expression increased irrespective of the presence of MMC, but was still sensitive to the addition of the arbitrium peptide (Fig. 2H). Our results therefore point to a mechanism of integration of arbitrium and DNA damage pathways by combined AimR activation and LexA repression of *aimX* transcription.

Finally, we searched and identified clade 1 *aimR* receptor genes in near 300 genomes from the *B. subtilis* group of species. All these receptors were part of ϕ106 type phages (methods). We found that all these phages had a conserved core structure of the lysogeny module with a CI-like protein in an antisense direction to the putative *aimX*. Yet, these genomes varied in the arbitrium peptide sequence and genomic organization of putative lysis genes directly downstream of the lysogeny module (Supplementary Fig. 11). Despite this variation, these phages shared the presence of a DNA sequence with strong similarity to the canonical LexA binding site at several hundred base-pairs downstream of the *aimP* gene (Fig. 2E, Supplementary Fig. 11). In summary, our results imply that in this phage family, the integration of DNA damage and communication is conserved and occurs at the *aimX* promoter.

### Prophage signaling alters infection dynamics

It was previously shown that infecting phages communicate with each other using AimP, and here we have demonstrated that dormant prophages communicate among themselves using this same signal. This opens up the possibility of communication between phages at different stages of their life cycle. Particularly, signaling by prophages may also impact infection dynamics by tipping the decision of phages infecting nearby permissive bacteria towards lysogeny. This stands in contrast to λ prophages, for example, which cannot directly impact the lysis-lysogeny decision of phages infecting neighboring cells.

To test the effect of signaling by prophages on the lysis\lysogeny decisions made by infecting phages, we grew co-cultures of 90% SPβ lysogens and 10% permissive, non-lysogenic, cells and infected them with WT phages (Fig. 3A, B). Lysogens were either Δ*aimR* mutants (which still produce the signal but are unable to induce) or Δ*aimRP* mutants (which neither produce the signal nor induce). If signaling by lysogens indeed affects the lysis\lysogeny decisions of infecting phages, phages infecting permissive bacteria grown with signaling lysogens are expected to choose lysogeny over lysis more frequently than phages infecting co-cultures composed of non-signaling lysogens. Co-cultures were grown in minimal medium so as to maximize the effect of signaling by prophages. Infecting phages, lysogens and non-lysogens were marked with different antibiotic resistance markers so that lysogens obtained from newly infecting phages could be discriminated from the rest of the population (Fig. 3B, Methods).

**Figure 3:**
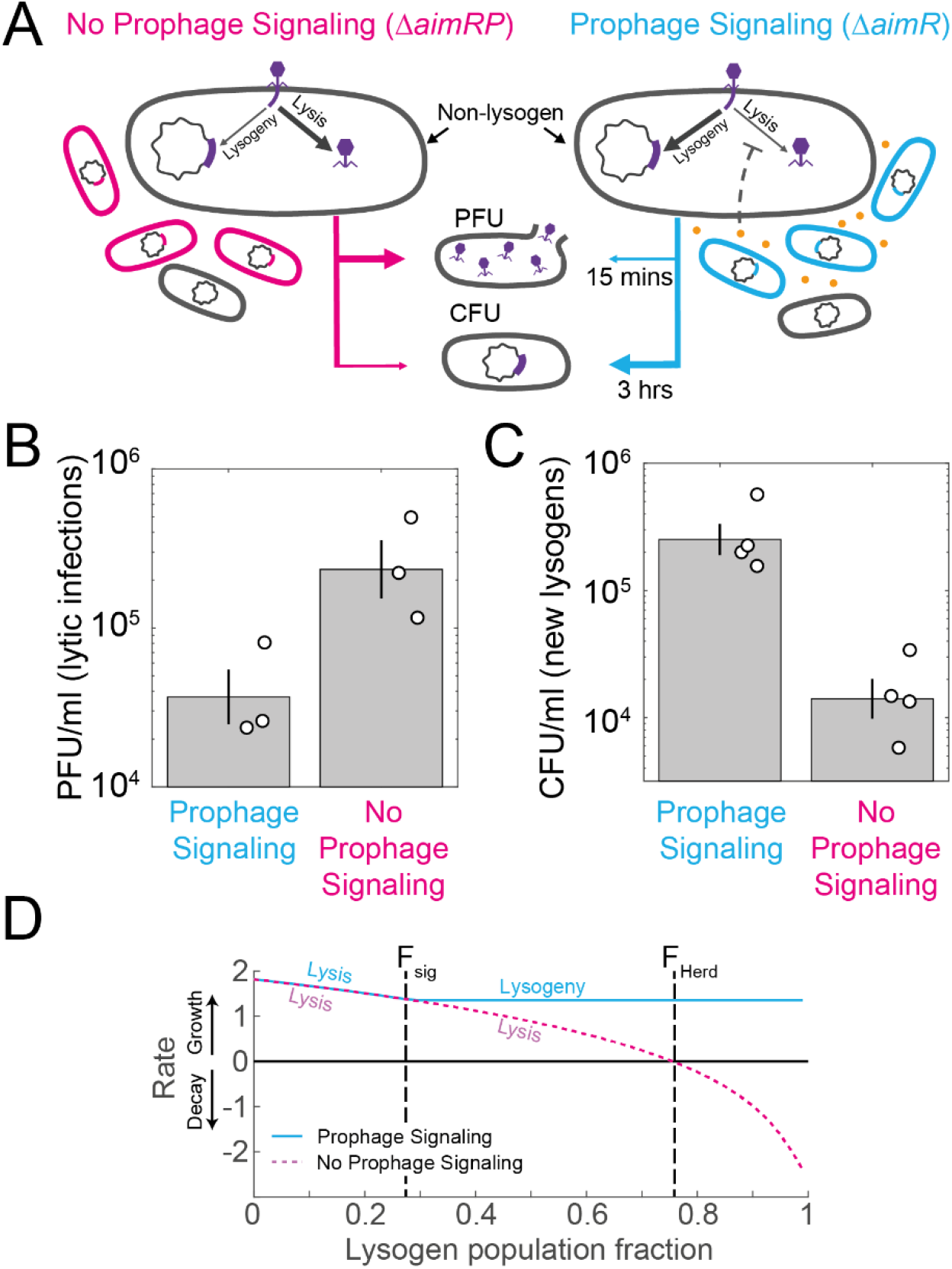
Prophage signaling biases infections towards lysogeny. (A-C) A scheme (A) and results (B, C) of the experiment. (A) A minority (10%) of permissive non-lysogens is incubated with either signaling (Δ*aimR*, cyan) or non-signaling (Δ*aimRP*, magenta) lysogens which do not produce phages. Cultures are infected by wild-type phages and monitored for PFU after 15 minutes (B) and for CFU after 3 hrs (C). Empty circles represent separate biological repeats and error bars represent standard errors. (D) A mathematical model of selection predicts an optimal threshold frequency for response to prophage signaling. Shown are the average rate of growth or decay of a phage infecting a permissive host as a function of the initial number of lysogens in the population (Supplementary Fig. 13). Two infection models are considered - with no prophage signaling (magenta) or with prophage signaling (cyan). The former is essentially lytic irrespective of lysogen fraction (Supplementary Fig. 12), while the latter switches from lysis to lysogeny above a signaling threshold corresponding to a population threshold *F*_*sig*_. Shown is the optimal threshold, where the switch occurs when lysogeny and lysis has the same growth rate. Also marked is the herd immunity frequency over which lytic infection is declining (*F*_*Herd*_). Details of the model are presented in Supplementary text.

We infected cultures at a total MOI of 0.1 and shortly after (15 minutes) performed a plaque assay without filtering out bacteria. This allowed us to track the lysis-lysogeny decisions made by phages during the first round of infections (Fig. 3A, methods). We found that the number of infecting phages which lysed bacteria and formed plaques was reduced by a factor of 6 in co-cultures with signal producing lysogens compared to those with signal null lysogens (Fig 3B, p=0.01, t-test for ratio of PFUs between cultures). Conversely, we found that the number of lysogens formed by infecting phages 3 hours after infection, was 17-fold higher in co-cultures containing signal-producing lysogens compared with signal null (Fig. 3C, p=0.02, t-test for ratio of CFUs between cultures). In total, the results show that under these conditions signaling by prophages indeed pushes infecting phages towards lysogeny, and by this protects surrounding bacteria from lysis (Fig. 3C).

### Modeling suggests that prophage signaling is beneficial

In this work, we showed that prophage signaling modulates both induction and lysis-lysogeny decisions, biasing phages towards lysogeny. Prophage signaling may thereby reduce lytic infection in the presence of lysogens in a manner unattainable by the co-infection-based lysis-lysogeny decision guiding phage λ among others, and provide an adaptive advantage to arbitrium-coding phages.

To better understand the impact of prophage signaling on phage decision making and population dynamics, we formalized two mathematical models of phages with different decision-making mechanisms upon infection – a λ-like, coinfection-sensing based mechanism, which cannot respond to the presence of lysogens, and an arbitrium-like mechanism which responds to the presence of lysogens through prophage signaling (See Supplementary Text for description of the models). We used these models to study the infection dynamics of a small population of free phage either in the absence of lysogens, or during initial infection of a population containing a fraction *F* of lysogens. Notably, we assume that lysogens are susceptible to phage adsorption, but protected from formation of a viable infection cycle, as is true for phage λ, for phage SPβ (Supplementary Fig. 3) and many others. We find that while phages had a similar infection dynamic with no initial lysogen population, they differ considerably in the presence of such population (Supplementary Fig. 12). To characterize this difference, we defined the total growthof the phages as the average growth rate over time of phages in all their forms - free, infecting or lysogenic - per an initial successful infection (i.e, phage infection of a permissive cell).

Under this scenario, λ-like phages became lytic due to the low density of free phages, irrespective of lysogen frequency. This led to positive phage growth for lysogen frequencies lower than the herd immunity frequency (*F*_*H*_ = 1 ‒ 1/*R*, where *R* is the basic reproductive number of the phage) and phage population demise above this frequency (Fig. 3D). For arbitrium-like phages, we assumed that transition to lysogeny occurs at a threshold level of signaling, corresponding to a lysogen fraction threshold (Supplementary text). We find that the optimal signaling threshold occurs at the frequency where lytic growth rate is equal to the lysogenic one (Fig. 3D), which is lower than the herd immunity frequency (Supplementary Fig. 13, Supplementary Text). Therefore, prophage signaling may allow phages to optimize their fitness by coordinating lysogenization when free phage density is low but lysogen density is high. However, if the signaling threshold is lower than optimal, a tradeoff occurs with an advantage to λ-like phages over a range of frequencies where the communicating phages switch to lysogeny prematurely (Supplementary Fig. 13). Altogether, this analysis points to the adaptive value of prophage signaling in the regulation of phage life-cycle transitions.

## Discussion

Several phenomena which may be considered communication among phages have been previously shown to affect the lysis-lysogeny decision upon infection. These include interactions between phages during co-infection^12,13,34,35^, as well as the newly discovered arbitrium peptide-based communication ^5,6^. Recently, prophages have also been shown to eavesdrop on the cell-cell communication of their bacterial host to control induction^36,37^. Many other phages are known to code for quorum-sensing systems or receptors with an unknown function (e.g. ^38–40^).

In this work, we showed that intercellular communication between prophages controls induction. We found that arbitrium systems of two different phage families continue to function in the prophage state and regulate induction in conjunction with DNA damage. In addition, signaling by prophages biases infecting phages towards lysogeny. Mathematical modeling indicates that prophage communication may be an adaptive strategy of phages aimed at reducing the risk of detrimental lysis of a host (either an established host, or a potential one) in an environment where lysogens are already prevalent. Prophage signaling, as opposed to communication exclusively during active infections, thus provides a benefit that cannot be gained by previously known mechanisms.

The mechanism by which *aimX* promotes initiation of the lytic cycle and its interaction with the DNA-damage detection network in SPβ-like phages remains unknown, stressing the need to further study the lysogeny system of this poorly characterized family ^42^. Nevertheless, we have shown that in these phages, the inputs of DNA damage and arbitrium signaling are integrated downstream of *aimX* expression. In other arbitrium designs, as the one we explored in the ϕ106 phage of *B. subtilis subsp. Inaquosorum*, it has been suggested that *aimX* directly inhibits the expression of the phage CI-like repressor by cis-antisense expression ^6^. We showed here that in this system the DNA damage SOS response factor LexA, directly regulates *aimX* in concert with AimR. The striking difference in the integration of the two sensory pathways, as well as other differences in the genetic organization between these arbitrium clades, may suggest that the integration of DNA damage and arbitrium signals evolved independently in the two phage families.

In nature, bacteria are thought to often reside in dense spatially structured communities^43^. Recent analysis suggests that under such conditions, RRNPP-type quorum-sensing systems report for the local frequency of signal producers rather than their overall density^22^. In a dense structured community, the local frequency of lysogens is similarly a better predictor of successful future infections than the makeup of the population more globally. Therefore, the use of short-range communication may be especially advantageous in a natural setting.

Additionally, the short range of arbitrium communication coupled with the clonal nature of bacterial populations at this local scale may also help relax the conditions under which prophage signaling is beneficial (Supplementary Fig. 13). Mathematical modelling indicates that the point at which the tradeoff posed by the strategy of lysogen signaling becomes beneficial, depends on several epidemiological parameters which may vary in natural conditions. If phages are likely to encounter communities which at a local scale are completely dominated by either lysogenic or non-lysogenic bacteria, responding to the existence of lysogen becomes significantly more robust with respect to these parameters (Supplementary text) ^22,44^.

In both phage families we explored, repression of induction by intercellular communication is dominant over the activation of induction by DNA damage. In contrast, some other mobile elements which rely on similar inputs have a different mode of integration. The induction of conjugation in ICE*Bs*1, a *B. subtilis* integrative and conjugative element, is also controlled by both DNA damage and quorum-sensing. Here too, the communication signal reduces the rate of induction under different conditions. However, in the case of ICE*B*s1 DNA damage alone is sufficient to induce conjugation, independently of the communication signal ^41^. A similar logic of integration was found in the regulation of induction by DNA damage and host signals in a *Vibrio* phage ^29,33^. It would be interesting to understand the evolutionary forces that drive these differences and their ecological consequences.

## Methods

### Strain construction

All bacterial strains, plasmids and primers used in this study are listed in supplementary Tables 1 - 3. To construct new *B. subtilis* strains, standard transformation and Spp1 transduction protocols were used for genomic integration and plasmid transformation^45^. The *B. subtilis* lab strain PY79 contains an additional DNA damage-induced lytic element (the PBSX prophage-derived bacteriocin) and therefore was only used for gene expression experiments ^27^. For plaque and growth assays we used a *Δxpf* strain in which PBSX cannot induce ^46^. Experiments with the arbitrium system of ϕ106 were conducted on the background of a PY79 strain where PBSX is deleted at its entirety, and ϕ3T experiments were conducted on Δ6 background which lacks all mobile elements ^18^.

Strain SPβ used in this work was originally described as wild-type by Erez et al ^5^. Notably, subsequent to the experiments described in this work, we have recently found through sequencing and perturbation experiments that the SPβ phage strain we used harbors mutations which render it heat-inducible (SPβc2 strain) ^47^. Importantly, we corroborated the effect of the arbitirum peptide on induction of wild-type, heat insensitive SPβ, showing that these mutations do not affect our observed phenotype (Supplementary Fig. 14).

*aimP, aimR* and *xpf* were deleted by transformation of kanamycin resistance cassettes from a *Bacillus subtilis* 168 deletion library^48^ into the appropriate strains. The kanamycin resistance cassette was excised from the *xpf* deletion using a Cre/*lox* system^49^. For the *aimRP* and PBSX deletions, we used a long flanking homology PCR method to replace both genes with a kanamycin resistance cassette^49^. All primers used are listed in supplementary table 3.

### Plasmid construction

pAEC1563 was constructed by amplifying the *aimX* promoter using the POL01 and POL02 primer pair. The PCR product was digested with BamHI-HF and NheI-HF, and cloned into pAEC277 digested with NheI-HF and BamHI-HF.

pAEC1909 was constructed by amplifying the arbitrium locus (AimR, AimP, aimX) using stav128 and stav132 primer pair from *B. subtilis* subsp. *inaquosorum* KCTC 13429 (AES7108) genomic DNA. The PCR product was digested with BamHI-HF and MfeI-HF, and cloned into pAEC277 digested with EcoRI-HF and BamHI-HF.

pAEC2079 was constructed by amplifying the whole pAEC1909 using stav152 and stav153 primer pair that contain the point mutation in the putative LexA binding site. The original plasmid in the PCR product was degraded using DpnI. The PCR product was closed after adding phosphate using PNK enzyme, followed by ligation.

pAEC2081 was constructed by amplifying lexAind from AIG246 ^24^ using the EM9 and EM10 primer pair. The PCR product was digested with SphI-HF and NheI-HF, and cloned into pAEC1505 digested with NheI-HF and SphI-HF.

pAEC1919 was constructed by amplifying the *dinC* promoter using the P200 and P201 primer pair. The PCR product was digested with BamHI-HF and NheI-HF, and cloned into pAEC277 digested with NheI-HF and BamHI-HF.

### Growth media and conditions

As a rich medium in this study, we used Lysogeny Broth (LB): 1% tryptone (Difco), 0.5% yeast extract (Difco), 0.5% NaCl. For experiments in minimal media we used Spizizen minimal medium (SMM; 2 g L^‒1^ (NH_4_)_2_SO_4_, 14 g L^‒1^K_2_HPO_4_, 6 g L^‒1^KH_2_PO_4_, 1 g L^‒1^ trisodium citrate, 0.2 g L^‒1^MgSO_4_ 7H_2_O), supplemented with trace elements (125 mg L^‒ 1^MgCl_2_ 6H_2_O, 5.5 mg L^‒1^CaCl_2_, 13.5 mg L^‒1^FeCl_2_ 6H_2_O, 1 mg L^‒1^MnCl_2_ 4H_2_O, 1.7 mg L^‒ 1^ZnCl_2_, 0.43 mg L^‒1^CuCl_2_ 4H_2_O,0.6 mg L^‒1^CoCl_2_ 6H_2_O, 0.6 mg L^‒1^Na_2_MoO_4_ 2H_2_O). 0.5% glucose served as a carbon source. Liquid cultures were grown with shaking at 220 RPM and a temperature of 37°. When preparing plates, medium was solidified by addition of 2% agar. Antibiotics were added (when necessary) at the following concentrations: spectinomycin: 100 μg ml^-1^, chloramphenicol: 5 μg ml^-1^, kanamycin: 10 μg ml^-1^, MLS: 3 μg ml^-1^ erythromycin + 25 μg ml^-1^ lincomycin. For experiments that involve infection, media were supplemented with 0.1 mM MnCl_2_ and 5 mM MgCl_2_.

### Flow cytometry

Flow cytometry was performed to quantify gene expression at the single-cell level, using a Beckman-Coulter Gallios flow-cytometer equipped with four lasers (405 nm, 488 nm co- linear with 561 nm, 638 nm). The emission filters used were: BFP – 450/50, YFP – 525/40, mCherry – 620/30. Constitutive mCherry and mTag2-BFP were used to distinguish between co-cultured cells. Median YFP levels were measured relative to a set voltage which was approximately set such that a value of 1.25 will be given to autofluorescence of strain PY79 in SMM medium.

### Tracking expression by fluorescent reporters

#### Phage SPβ

To determine expression levels of *aimX* in SPβ lysogens grown in rich media, strains harboring an *aimX*-YFP reporter were grown in LB overnight at 37° with shaking at 220 RPM, and then diluted by a factor of 1:100 into fresh LB media containing 10 μM of the SPβ arbitrium peptide (GMPRGA) when indicated. Upon reaching OD_600_ = 0.3, fluorescence was measured using flow cytometry. To measure expression in co-cultures, after reaching OD_600_ = 0.3, strains were mixed at a 50:50 ratio. Co-cultures were grown ON and subsequently diluted, regrown and measured for fluorescence in the same way as mono-cultures.

To track gene expression in response to mitomycin C, strains harboring fluorescent reporters for the transcription of *aimR* or *aimX* were grown in LB overnight at 37° with shaking at 220 RPM, and then diluted by a factor of 1:100 into fresh LB media. Upon reaching OD_600_ = 0.3, 0.5 μg ml^-1^ of mitomycin C (Sigma, M4287; MMC) was added to the medium, and both YFP and BFP were measured every 15 minutes using flow cytometry. To determine expression in minimal media, strains were grown to OD_600_ = 0.1 in SMM containing trace elements and glucose at 37° with shaking at 220 RPM. At this stage strains were mixed at different frequencies as indicated. Cultures were then diluted by a factor of 10^4^-10^6^ into fresh SMM media supplemented with 10 μM of the SPβ arbitrium peptide when indicated, and grown for about 12 hours in exponential phase. Optical density and gene expression were measured at approximately 1-hour intervals using a spectrophotometer and a flow cytometer respectively. For very low densities, optical density at time of measurement was estimated by extrapolating backwards from optical density at later times assuming a growth rate of 50±8.5 minutes supported by growth rate measurements, where we fitted growth data (OD_600_ vs time) to an exponent.

#### Phage ϕ106

Strains harboring the arbitrium system of phage ϕ106 upstream of a YFP fluorescent reporter and a constitutive BFP reporter at a different locus (see plasmid construction and tables 1-3) were used determine *aimX* expression in this system under different conditions. Cultures were grown in LB overnight at 37° with shaking at 220 RPM, and then diluted by a factor of 1:1000 into fresh LB media containing different concentrations of IPTG. Upon reaching OD_600_ = 0.3, 10 μM of the appropriate arbitrium peptide (DPPVGM) and/or 0.5 μg ml^-1^ of MMC were added when indicated (t=0). YFP and BFP were measured at several time points.

### Plaque forming assay

Samples for PFU measurements were collected from cultures centrifuged for 5 min at 4,000 r.p.m at room temperature. Next, the supernatant was filtered using a 0.2 μm filter (Sartorius Stedim biotech cat. 14-555-270). 100 μL of filtered supernatant at an appropriate dilution was then mixed with 200 μL of *B. subtilis* PY79 grown to OD_600_ in MMB (LB supplemented with 0.1 mM MnCl_2_ and 5 mM MgCl_2_), and left to incubate at room temperature for 5 minutes. Three ml of molten LB-0.5% agar medium (at 60°C) supplemented with 0.1 mM MnCl_2_ and 5 mM MgCl_2_ was added, mixed, and then quickly overlaid on LB-agar plates. Plates were then incubated at 37° for 1 hour and then ON at room temperature to allow plaques to form.

### Induction rate

For induction in rich medium, strains were grown overnight in at 37° with shaking at 220 RPM, and then diluted by a factor of 10^4^ into fresh LB media containing 10 μM of peptide when indicated. To assess levels of spontaneous induction, samples for PFU measurements were taken at OD_600_ = 0.3. To measure induction in the presence of DNA damage, 0.5 μg ml^-1^ of MMC was introduced at OD_600_ = 0.3 and samples for PFU measurements were collected after 120 minutes.

For induction in minimal medium strains were grown to OD_600_ = 0.1 at which point cultures were mixed as indicated, and diluted by a factor of 10^6^ in fresh SMM media. To assess spontaneous induction, samples for PFU measurements were taken at OD_600_ = 0.3. To measure induction in the presence of DNA damage, MMC was introduced at OD_600_ = 0.3 and samples for PFU were collected after 150 minutes.

### Induction growth dynamics

To examine growth dynamics during prophage induction, strains were grown overnight in LB at 37° with shaking at 220 RPM then diluted by a factor of 1:100 into fresh LB media. Upon reaching OD_600_ = 0.3 cultures were supplemented with 0.5 μg ml^-1^ of MMC and 10 μM of peptides when indicated. Optical density measurements at a wavelength of 600 nm were performed in a 96-well plate using a plate reader (PerkinElmer VICTOR Nivo Multimode Plate Reader).

### Infection experiments

Strains were grown at 37° with shaking at 220 RPM to OD_600_ = 0.1 in SMM containing trace elements and glucose and then mixed for co-cultures at the indicated frequencies. Cultures were then diluted by a factor of 10^6^ into fresh SMM media containing 0.1 mM MnCl_2_ and 5 mM MgCl_2_, grown until reaching OD_600_ = 0.3 and infected with free SPβ phages harboring a chloramphenicol resistance marker at an MOI of 0.1. Samples were taken after 15 minutes without filtering, and mixed with a phage-free indicator strain in order to perform plaque assay. Additional samples were taken 3 hours after the time of infection to measure CFU on plates of agar supplemented with chloramphenicol.

### Real Time qPCR Measurements

Total RNA was extracted from cells using a High Pure RNA Isolation kit (Roche). To this end, cells were grown in LB at 37° with shaking at 220 RPM to OD_600_ = 0.3 at which point MMC was added along with the arbitrium peptide when indicated. Samples of different timepoints were centrifuged for 5 minutes at 2000 RCF and pellets were flash frozen in liquid nitrogen. 1 μg of RNA was reverse-transcribed to cDNA using a qScript™ cDNA Synthesis Kit (Quanta BioSciences). Real-time qPCR was performed on a Step One Plus Real Time PCR system (Applied Biosystems), using SYBR Green (Quanta BioSciences). Specificity of primers (listed in table 3) was validated using a melt curve. Efficiency of primers used was between 95% and 105%, with calibration curves of r^2^ > 0.95. Thermocycling parameters were as follows – Holding stage at 95° for 1 minute, and 40 cycles of 2 steps – a first step at 95° for 5 seconds and a second step at 60° for 30 seconds. Transcript levels were normalized to levels of the reference gene *rpoB*. RNA samples were stored at -80° and cDNA was stored at -20°. Nucleic acid quantification was performed by Thermo Scientific NANODROP 2000c Spectrophotometer. Results were analyzed using the Step One™ v2.3 software by the standard ΔΔCt method. Each condition was measured across three biological repeats, where each biological repeat included two technical repeats. In general, unless mentioned otherwise reagent concentrations and volume, time, temperature, and general conditions used were as detailed in the accompanying kit protocol.

### Adsorption experiments

Strains were grown at 37° with shaking at 220 RPM to OD_600_ = 0.1 in SMM containing trace elements and glucose and then diluted by a factor of 10^6^ into fresh SMM media containing 0.1 mM MnCl_2_ and 5 mM MgCl_2_, grown until reaching OD_600_ = 0.3 and infected with ∼10^6^ SPβ phages. Samples were taken after 15 minutes, centrifuged, and filtered using a 0.2 μm filter (Sartorius Stedim biotech cat. 14-555-270).. Free phages remaining in the medium were then enumerated by plaque assay.

### Phylogenetic tree reconstruction, SOS motif discovery and comparison between phages

We collected 297 genomes of *B. subtilis* group genomes. We performed a BlastP search for all AimR homologs in these genomes using a query based on reprasentatives of all nine AimR clades^6^, yielding a total of 401 variants. Of these, 81 clustered with AimR of *B. subtilis subsp. Inaquosorum* ϕ106 strain and clade 1 AimRs, while the rest belonged to clades #2,3. We identified the *aimP* gene and its putative mature AimP signal, as well as other nearby open reading frames for all of these strains. We also verified by blast the existence of a repressor homolog in an antisense direction to the *aimRP* operon. All 81 variants had a LexA binding site of high similarity to the consensus sequence between *aimR* and the repressor. LexA binding site was identified using the algorithm provided by the database DBTBS ^33^. Nine representative sequences were chosen, which have a varying AimP peptide or gene organization, including the one from *B. subtilis subsp. Inaquosorum*. These were used to create Supplementary Fig. 11. NCBI accession numbers for these genes are given in Supplementary Fig. 11 legend. Comparison between phages ϕ106 and ϕ105 was done by BlastP for all open reading frames as described in the legend of Supplementary Fig. 10.

## Notes

### Competing Interest Statement

The authors have declared no competing interest.

