## supplements for "Dormant phages communicate to control exit from lysogeny"

Infection dynamics for coinfection and arbitrium-like sensing mechanisms are similar. 30

Early infection in the presence of lysogens 30

Adaptivity of signaling in structured populations 31

References .......... 33

#### Supplementary Tables

##### Table S1: Strain list

| Strain name | Background | Genotype | Source (ref) | Strain designation (used in Figure) |
|---|---|---|---|---|
| AES2837 | PY79 | *Bacillus subtilis* PY79, wild type | Lab stocks | Used as indicator strain for plaque assays |
| AES6534 | PY79 | *amyE*::(P*aimX*-*aimX*-3xYFP-Spec) | This work | Non-lysogen (1A) |
| AES6533 | PY79 | SPβ c2 lysogen *amyE*::(P*aimX*-*aimX*-3xYFP-Spec) | This work | wt SPbeta (1A) |
| AES6764 | PY79 | SPβ c2 lysogen Δ*aimR*:Kan; *amyE*::(P*aimX*-*aimX*-3xYFP-Spec) | This work (Δ*aimR* transferred from mutant library [1]) | Δ*aimR* (1A) |
| AES6242 | PY79 | SPβ c2 lysogen Δ*aimP*:Kan; *amyE*::(P*aimX*-*aimX*-3xYFP-Spec); *lacA*::(P*veg*-R0-2xmTag-BFP-Mls) | This work | Δ*aimP* (1A, B, S2, S4) |
| AES7404 | PY79 | Δ*xpf* *amyE*::(P*aimX*-*aimX*-3xYFP-Spec) | This work | S5 |
| AES6969 | PY79 | Δ*xpf* | This work (Δ*xpf* transferred from mutant library [1]) | Non-lysogen (1C, S3) |
| AES6904 | PY79 | SPβ c2 lysogen Δ*xpf* | This work | wt (1C, D, S1) |
| AES6939 | PY79 | SPβ c2 lysogen Δ*xpf*; Δ*aimP*::Kan | This work (Δ*aimP* transferred from mutant library [1]) | Δ*aimP* (1C, D, S1) |
| AES7265 | PY79 | SPβ c2 lysogen Δ*xpf*; Δ*aimR*::Kan | This work (Δ*aimR* transferred from mutant library [1]) | Δ*aimR* (1C, S1, S3) |
| AES6549 | PY79 | SPβ c2 lysogen *amyE*::(P*aimX*-*aimX*-3xYFP-Spec); *lacA*::(P*veg*-R0-mCherry-Mls) | This work | wt (1B) |
| AES6551 | PY79 | SPβ c2 lysogen; *amyE*::(P*aimX*-*aimX*-3xYFP-Spec); *lacA*::(P*veg*-R0-mTag-BFP-Mls) | This work | P*aimX*-*aimX*-YFP (2B, S4) |
| AES5666 | PY79 | SPβ c2 lysogen *amyE*::(P*aimR*-3xYFP-Spec) | This work | P*aimR*-YFP (S4) |
| AES7772 | PY79 | *lacA*::(P*veg*-R0-mTag-BFP-Mls) *amyE*::(P*dinC* -3xYFP) | This work | P*dinC*-YFP (2B) |
| AES6143 | PY79 | SPβ c2 lysogen | This work | 2A |

| | | | | |
|---|---|---|---|---|
| AES7108 | B. subtilis inaquosorum KCTC 13429 | | from Rotem Sorek's lab | 2D |
| AES7021 | PY79 | Δ*xpf sacA*::(P*veg*-R0-mCherry-Cm) | This work | Non-lysogen (3B, C) |
| AES7275 | PY79 | SPβ c2 lysogen Δ*xpf* Δ*aimR*::Kan; *amyE*::(P*veg*(+1/+8)-R0-2xmTag-BFP-Spec) | This work | Signaling lysogen (3B, C) |
| AES7277 | PY79 | SPβ c2 lysogen Δ*xpf*; Δ*aimRP*::Kan; *amyE*::(P*veg*(+1/+8)-R0-2xmTag-BFP-Spec) | This work | Non-signaling lysogen (3B, C) |
| AES6360 | Δ6 | ϕ3T lysogen Δ*pks*:Cm | (168 cured of all mobile elements Dpks:Cm) From BGSC[2] infected with ϕ3T | ϕ3T (S9) |
| AES7552 | PY79 | ΔPBSX *amyE*::(*aimRPX*$_{\phi 106}$-3x-YFP-Spec); *lacA*::(P*veg*-R0-mTagBFP-Mls) | This work | 2F, S7B, S8A |
| AES7903 | PY79 | ΔPBSX *amyE*::(*aimRPX*$_{\phi 106}$-3x-YFP-Spec); *lacA*::(P*veg*-R0-mTagBFP-Mls) *yhdHG*::(Phs-*lexA* ind-*lacI*-Kan) | This work | 2G, S7C, S8B |
| AES7905 | PY79 | ΔPBSX *amyE*::(*aimRPX*$_{\phi 106}$-3x-YFP-Spec) with mutated LexA binding site; *lacA*::(P*veg*-R0-mTagBFP-Mls); | This work | 2H, S8D |
| AES7913 | PY79 | Δ*xpf yhdHG*::(Phs-*lexAind*-*lacI*-Kan) *amyE*::(P*dinC*-YFPx3-Spec) | This work | S8C |
| AES7975 | PY79 | Δ*xpf* SPβ lysogen | This work | Heat insensitive (S14) |

#### Table S2: Plasmid list

| **Name** | **Description** | **Reference** |
|---|---|---|
| pAEC277 | pDL30::3xYFP-Spec (Amp) | Lab stock |
| pAEC1563 | pDL30::P*aimX*-*aimX*-3xYFP (Amp) | This work |

| | | |
|---|---|---|
| pAEC1909 | pDL30-*aimRPX*$_{\phi106}$-3x-YFP (Amp) | This work |
| pAEC2079 | pDL30-*aimRPX*$_{\phi106}$-YFP; LexA mutated binding site (Amp) | This work |
| pAEC1505 | pMMH253:Phs-*lacI* (Amp, Kan) | Lab stock |
| pAEC2081 | pMMH253-Phs-*lexA ind*-*lacI* (Amp) | This work |
| pAEC1919 | pDL30-P*dinC*-YFPx3 (Amp) | This work |

**Table S3: Primer list**

| primer name | sequence | used for |
|---|---|---|
| stav106 | TCAGGCGTTTCAATCGGACA | RT *rpoB* (reference gene) fw |
| stav107 | GCCGTCTGTCAGCATTAGGA | RT *rpoB* (reference gene) rev |
| stav108 | CCAAATGTCAGGACCCCGTT | RT holin fw |
| stav109 | GCTTGGTTCGGTTATGTGCG | RT holin rev |
| stav112 | GCCCGCTTCCTTGTGATTTC | RT replication related gene fw |
| stav113 | TGCGTCCTGAAACATTGTTCG | RT replication related gene rev |
| stav114 | CCACTGGAACATTTAACACTTGAG | RT putative early gene fw |
| stav115 | GCTTTCTCGCTCCTATATTAGTAAG | RT putative early gene rev |
| stav128 | GTACCAATTGGGAAAAAATCCACAAAGTATGATACTAAGG | Arbitrium locus phi105-like fw |
| stav132 | ATGAGGATCCCTGACTCATTCTCTAATAACTGCTTC | Arbitrium locus phi105-like rev |
| stav152 | CCTTTTGCGTTCGTTTGTTCTTATTATAATTG | LexA binding site point |

| | | |
|---|---|---|
| | | mutation p1 |
| stav153 | GGCTACTAACAATGAAAAAAGAAATTTCAC | LexA binding site point mutation p2 |
| POL127 | TCGTTACCTTGGCATTCACA | RT *rpoB* fw |
| POL128 | CACGGTTATCAAACGGCTCT | RT *rpoB2* rev |
| POL131 | GGAACCTTACCAACGTTATGAGG | RT *sprB* fw |
| POL132 | GTCATTACTTGCTTCATCCTGAGT | RT *sprB* rev |
| POL133 | TGGTTTTGATGCATTCGGGG | RT *yorB* fw |
| POL134 | CCATCGTCTTTAGCCCCTTC | RT *yorB* rev |
| POL135 | CGCCCTGACAAACTGTTGATTG | RT *yonH* fw |
| POL136 | GGTCAAATCACCACTGGGAGAC | RT *yonH* rev |
| POL137 | GCATTTGCTGCTTTGGGATGG | RT *cwlP* fw |
| POL138 | CATTGGTTTGCTGGCTCTGC | RT *cwlP* rev |
| SOB369 | ACGGTAGAGAGAGCACAGATACGGCGGCATAAAATC CATTGACACATAAAGT | Δ*aimRP*-P3 |
| SOB370 | GAACGGTATCCTGCCTTTCCTCCCTCGCTATCCTTATTA ACTCCATTATTTCCC | Δ*aimRP*-P2 |
| SOB371 | CCGATAATGATGGTTGGGGAAC | Δ*aimRP*-P1 |
| SOB372 | GCATATCAATCATTGTCTCCCTC | Δ*aimRP*-P4 |
| POL01 | CGTCGGATCCCCAACAAGCTTCAGGTGATT | P*aimX* fw |
| POL02 | GGCGCTAGCGCATTTCATTTAATTAGGGTG | P*aimX* rev |
| SOB216 | GAGGGAGGAAAGGCAGGATACC | BKK fw |
| SOB217 | CGCCGTATCTGTGCTCTCTCTA | BKK rev |
| EM9 | CGATGCTAGCGAGGTGCGAAAAATGACGAAGCTATC A | NheI-*lexA*-F |
| EM10 | CGATGCATGCGGAAGAGGGGGTTATTTTATGCATCT G | SphI-*lexA*-R |
| P200 | CTGAGGATCCATGATGACACTTGTTCAAAACAG | P*dinC* BamHI |
| P201 | GTCAGCTAGCAAATTTACCACCTCTATTATTATTTG | P*din*C NheI |
| POL121 | GTTCTGCCCTATATAATCTCGG | ΔPBSX-P1 |
| POL122 | GAACGGTATCCTGCCTTTCCTCCCTCCGCCTCAGAAAC CAAATTTTTC | ΔPBSX-P2 |
| POL123 | ACGGTAGAGAGAGCACAGATACGGCGGACCATAAAA ATCCCGGAGC | ΔPBSX-P3 |
| POL124 | CTGGACATATGAGCCCAC | ΔPBSX-P4 |

| | | |
|---|---|---|
| POL125 | GCAGAAGATGTTTGTCAGTGC | ΔPBSX-P5 (VER) |

#### Supplementary figures

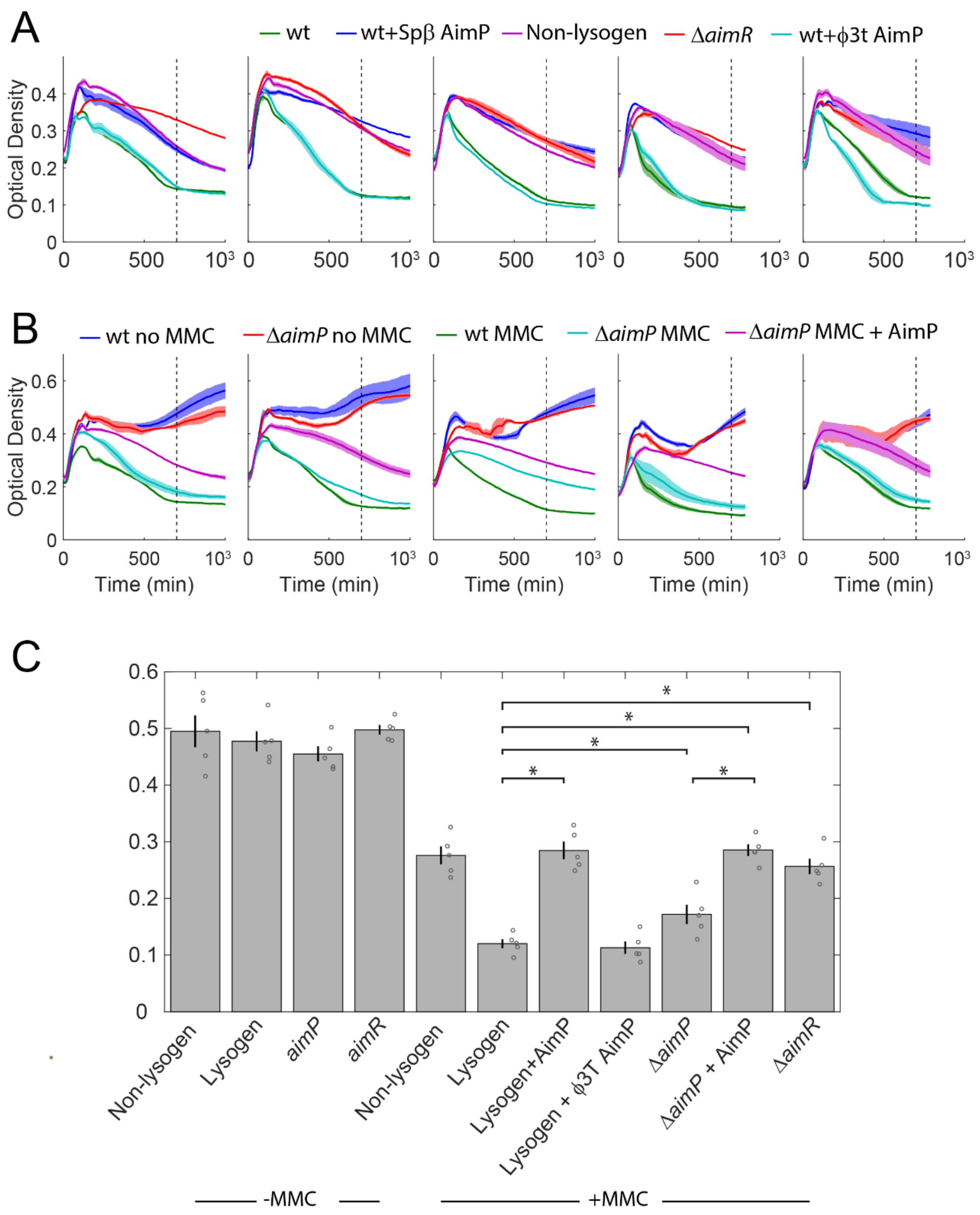

**Supplementary Figure 1: Impact of signaling on cellular growth curves as measured by plate reader.** (A) Further examples for plate reader experiments showing the same strains and conditions as shown in Fig. 1C of the main text. The second panel is the same as the panel shown in Fig. 1C. A dashed line marks the time-point at which OD levels were used for the statistical analysis shown in (C). (B) Plate reader experiments of additional strains and conditions. Shown are growth curves for the wild-type and

$\Delta$*aimP* lysogens with and without MMC,and for the $\Delta$*aimP* lysogen with MMC and AimP. See legend for the color of the curve of each strain. (C) Shown are the mean and standard error for each strain and condition, as well as individual measurements of optical density at T=700 minutes. Asterisks mark statistical significance ($p<0.05$, paired t-test).  Note that all differences between strains without MMC are non-significant.

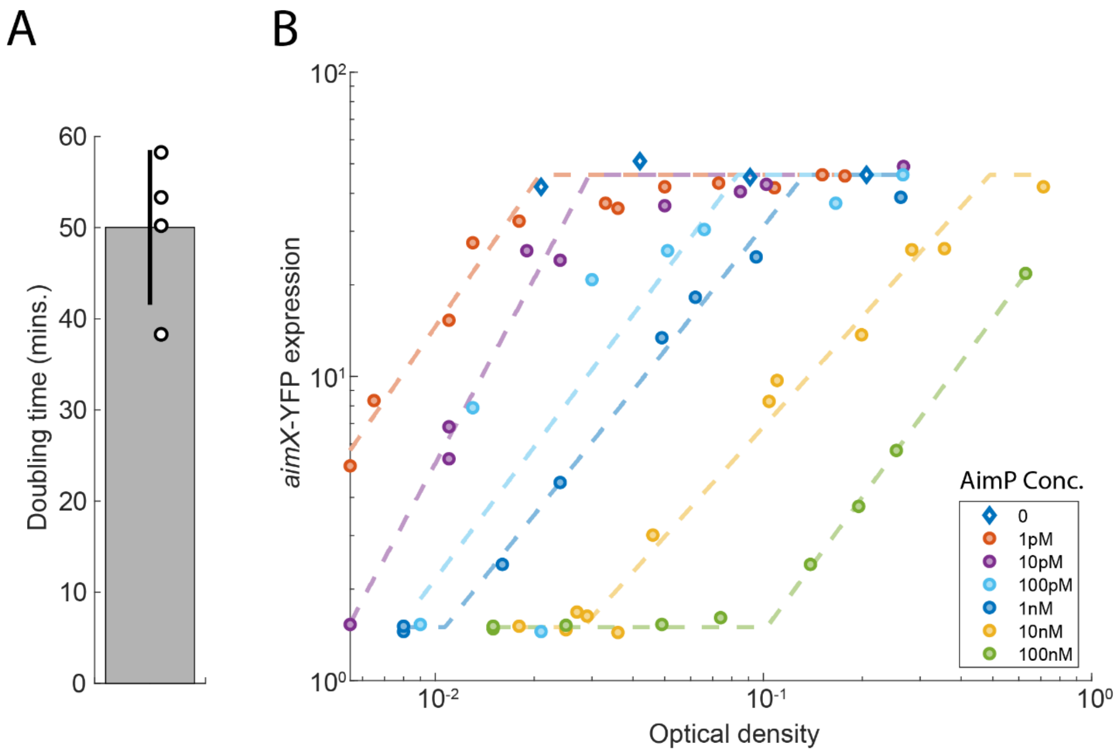

**Supplementary Figure 2: Sensitivity of the *aimX*-YFP reporter to addition of AimP in SPβ lysogens.** (A) Doubling time in minimal medium was measured by fitting the growth curve to an exponent. (B) Lysogens coding for the *aimX*-YFP reporter were diluted $10^6$-fold and grown overnight with the indicated levels of externally supplemented AimP peptide. Expression level and optical density were measured at different times. For low optical density, optical density was back-extrapolated from optical densities at later times and from the time passed by using a doubling time of 50 minutes (see A, as in Fig. 1B). Dashed lines show a tri-linear fit on the log-log scale of the data to two horizontal lines and an increasing line. The y-value of the horizontal lines of all fits is equal to the autofluorescence level (bottom line) and to the mean expression of cells with no signal (marked by diamond markers). The slope and intersect of the monotonously increasing lines are subject to fitting.

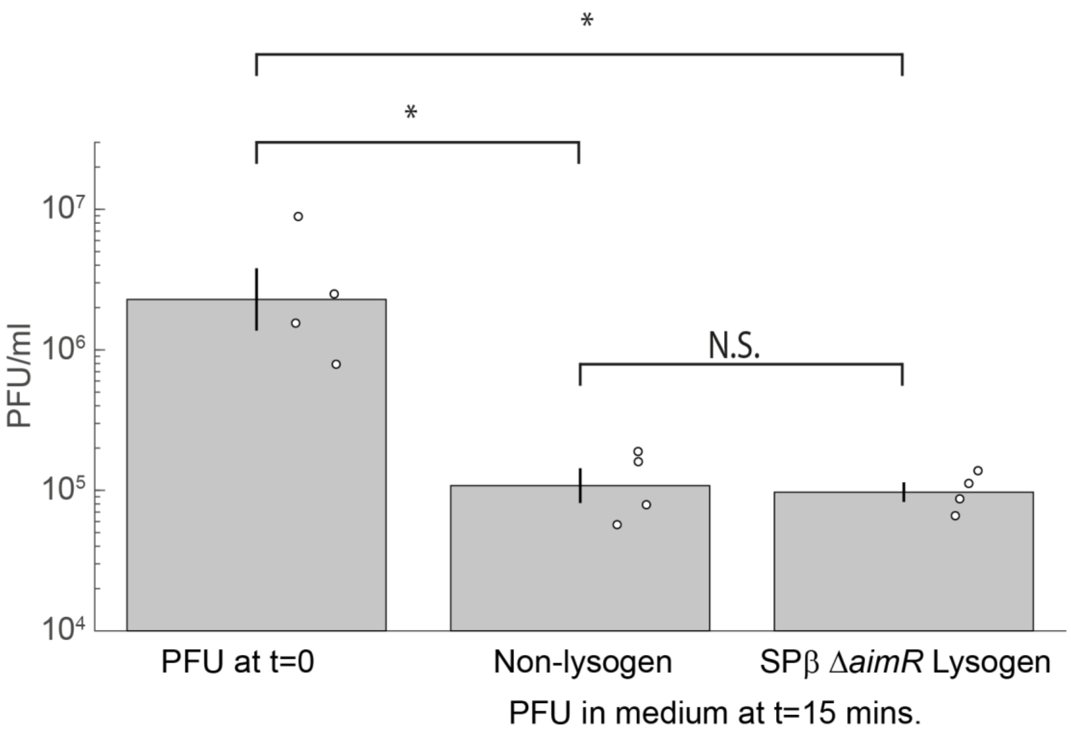

**Supplementary figure 3: SPβ adsorption to lysogens and non-lysogens is equal**. Shown are levels of SPβ phages, measured as PFU/ml, in the medium right after phage addition to medium with no cells and 15 minutes after addition of the same phage doses to medium containing either Non-lysogens or Δ*aimR* SPβ lysogens. The level of PFU is significantly lower in medium containing cells by a factor of ~30 ($p \leq 0.002$ for both cell types, paired t-test on the log of the PFU), but is not significantly different between the two cell types ($p=0.75$, paired t-test on the log of the PFU).

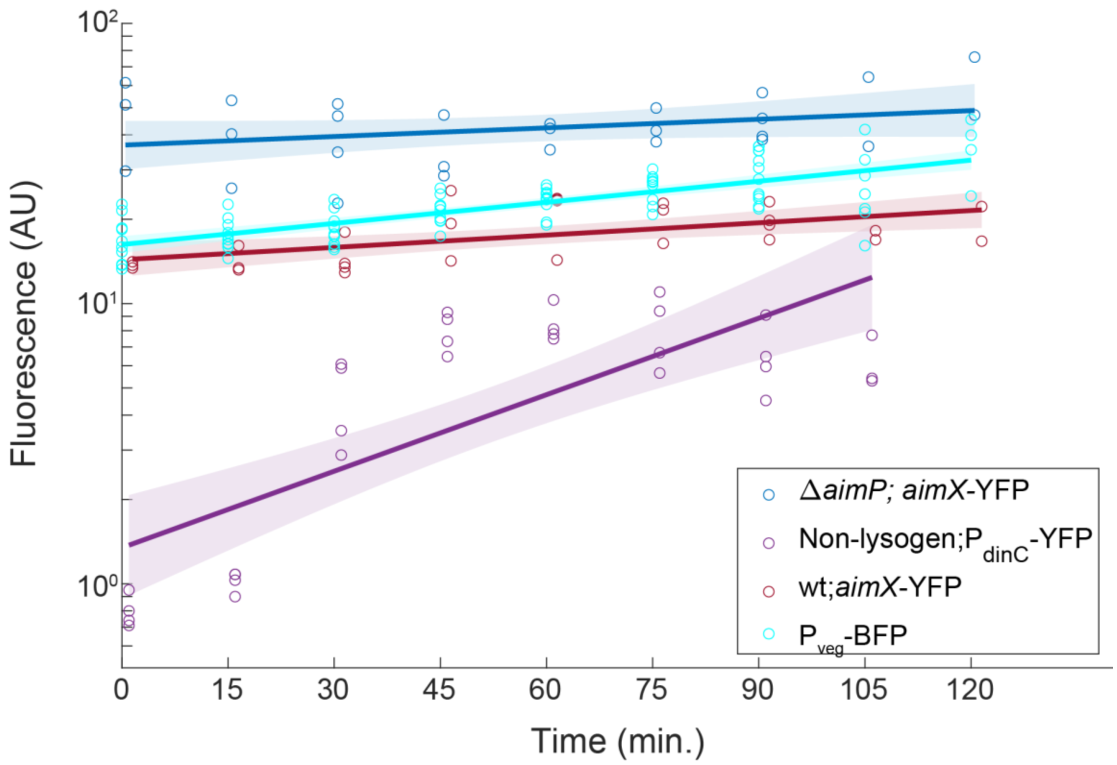

**Supplementary figure 4: YFP and BFP expression upon addition of MMC.** Shown are fluorescence levels (on a log scale) for different reporters and strains as a function of time from addition of MMC. An *aimX*-YFP reporter in wild-type (deep red) and in Δ*aimP* (blue) lysogens, a $P_{dinC}$-YFP reporter in a non lysogen strain containing PBSX (purple), and a $P_{veg}$-BFP constitutive reporter [3] plotted on a single curve for all the strains (cyan). Individual points are biological repeats, solid lines are linear best fit (with y data on a $\log_{10}$ scale) and shaded area is the boundary of error.

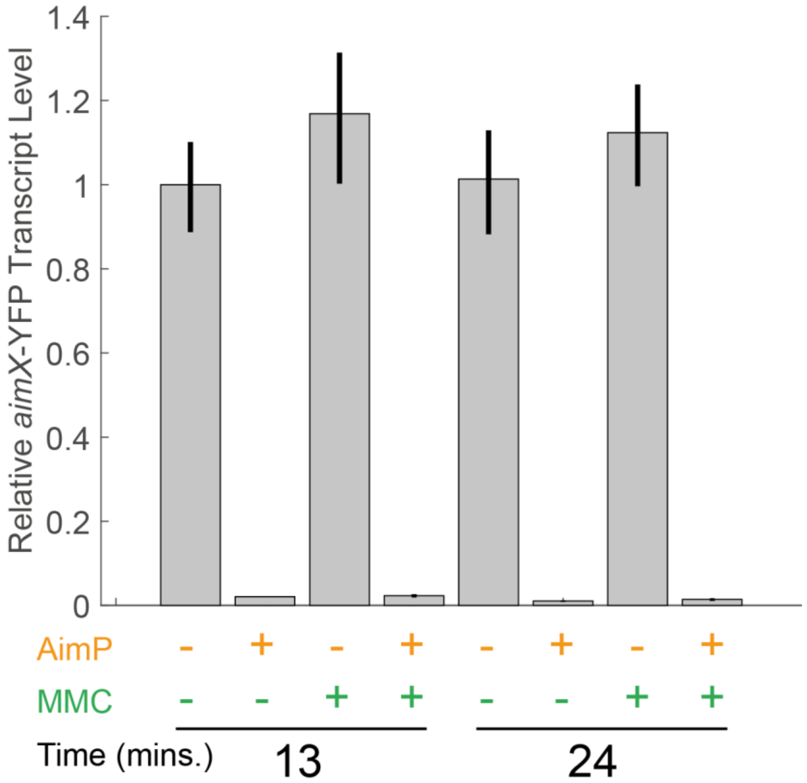

**Supplementary figure 5: RT-PCR of *aimX*-YFP under different conditions.** A Δa*imP* SPβ lysogen carrying the *aimX*-YFP reporter was assayed for its expression using RT-PCR (methods), either with or without the addition of 0.5μg/ml MMC and 10μM SPβ AimP. Measurements were taken either 13 or 24 minutes after the addition of MMC/AimP/Mock. Addition of MMC does not significantly alter *aimX*-YFP expression levels.

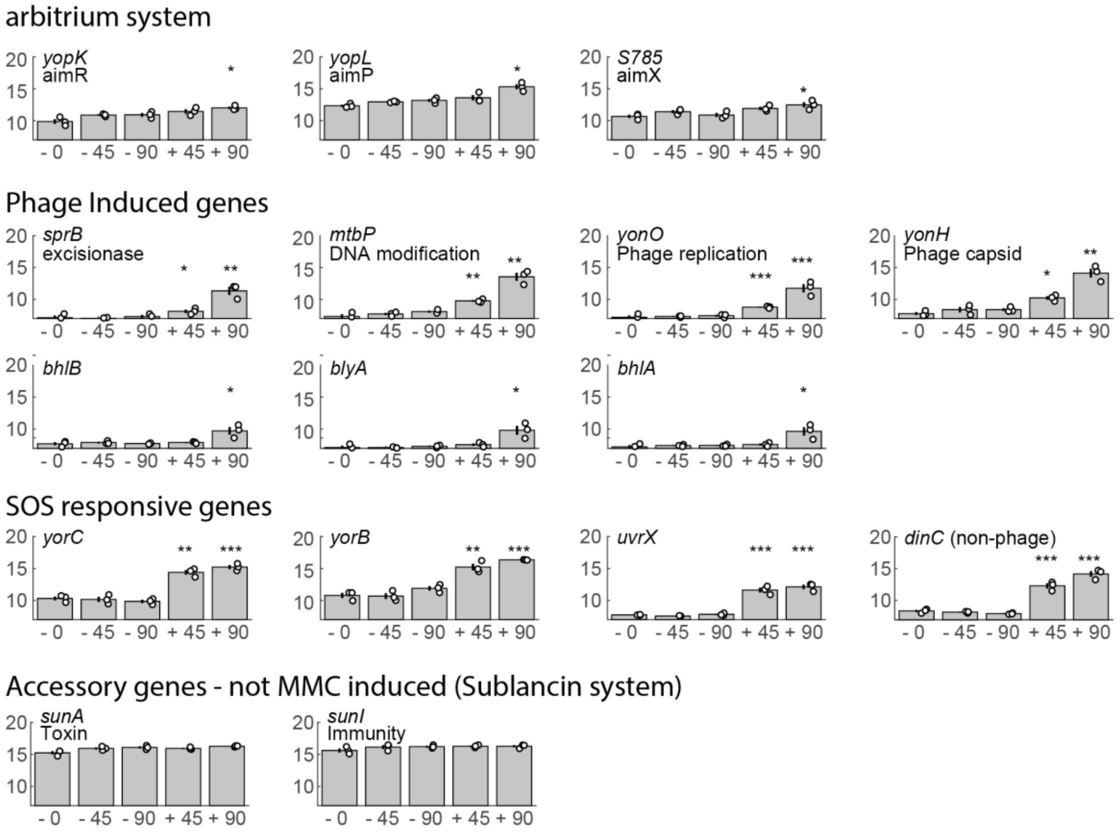

**Supplementary figure 6: Analysis of public RNAseq data of strain 168, after addition of mitomycin C (MMC)**. Shown are the RNAseq expression data from [4], for selected phage genes (and one non-phage gene, *dinC*) for 5 conditions – t=0 minutes (-0), t=45 minutes no MMC (-45), t=90 minutes no MMC (-90), t=45 minutes after MMC addition (+45), t=90 minutes after MMC addition (+90). Genes are divided into flour main categories based on their functional dependence. Gene name in the expression database and its current name or function are annotated. Marked with asterisks are cases where there is a statistically significant difference in the mean expression value with MMC compared to the corresponding time without it (* $0.01<p<0.05$; ** $0.001<p<0.01$; *** $p<0.001$, t-test comparison between the two cases). Specifically, note that *aimX*, *aimR* and *aimP* are not significantly different from the control 45 minutes after MMC addition and are marginally higher at 90 minutes. Note that strain 168 carries the mobile element ICEBs1 and contains several additional differences from the PY79 strain background we use in this work [5].

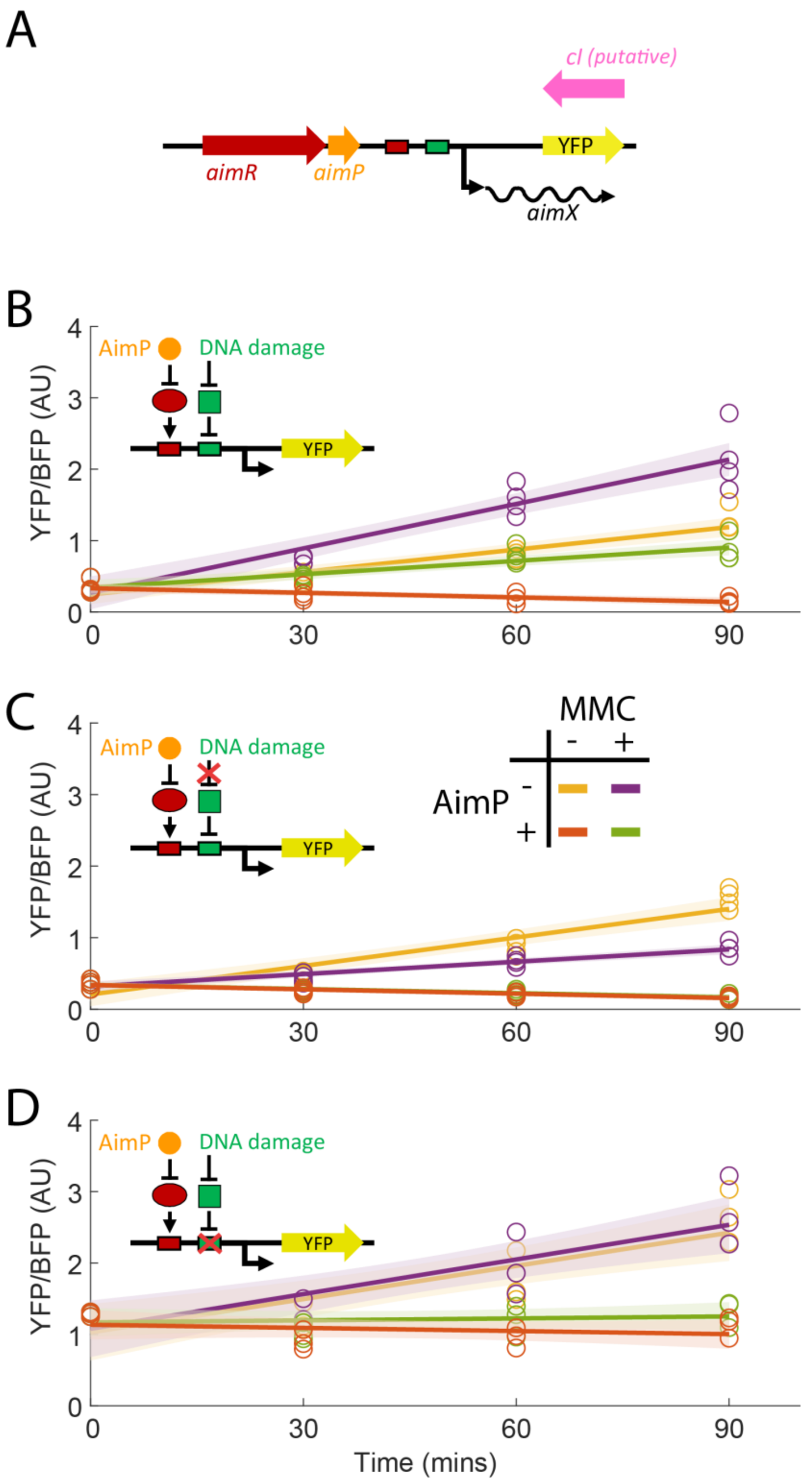

**Supplementary figure 7: *aimRPX*$_{\phi106}$-YFP construct and its expression as a function of time in three relevant genetic background.** (A) schematic structure of the *aimRPX*$_{\phi106}$-YFP reporter. The YFP genes starts where the putative cI gene ends in the native genome of the phage. (B-D) YFP expression from the *aimRPX*$_{\phi106}$-YFP construct as a function of time after addition of MMC and/or AimP (or mock additions), divided by fluorescence of a $P_{veg}$-BFP constitutive reporter within the same cells. Circles represent individual measurements (median YFP level from a flow cytometry measurement). Lines of the same color show linear best fit and shaded area show the boundary of error. Each strain was measured with three or four independent time series taken on different days. The legend in (C) is true for all panels. The three panels correspond to the same strains described in Fig. 2F-H of the main work and the illustrations describing them are identical; (B) *aimRPX*$_{\phi 106}$-YFP reporter based on the wild-type sequence of phage $\phi$106. (C) as in (A) but with overexpression of a non-cleavable defective lexA mutant. (D) a mutant *aimRPX*$_{\phi 106}$-YFP reporter with substitution in two base-pairs within the LexA binding site. Fig. 2F-H shows the data presented here for 90 minutes.

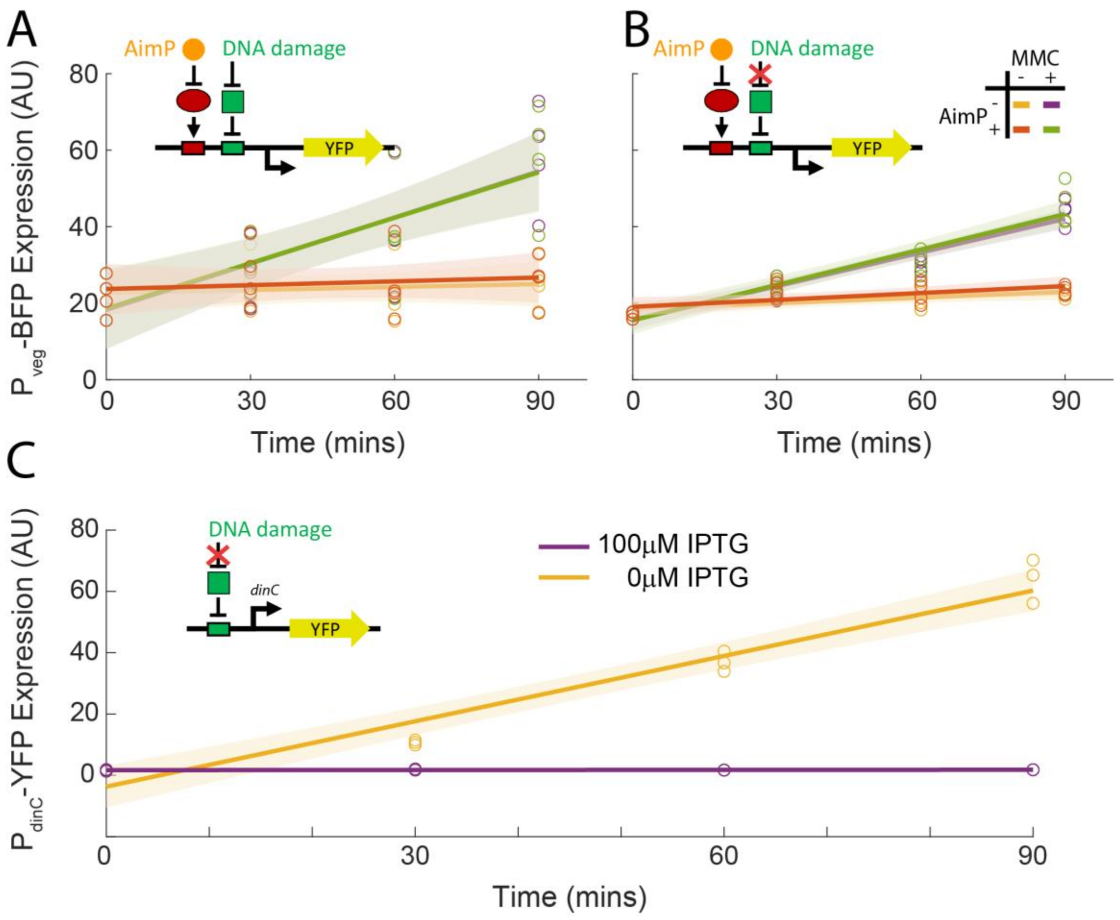

**Supplementary figure 8: $P_{veg}$-BFP and $P_{dinC}$-YFP expression upon addition of MMC with and without overexpression of a non-cleavable lexA mutant.** (A,B) Expression as a function of time of a constitutive **$P_{veg}$-BFP** reporter for the four conditions shown also in Supplementary Fig. 7. The two strains are the ones presented in panels A,B of Supplementary Fig. 7 correspondingly. Note the change in BFP expression upon addition in MMC between panels A,B, indicating the impact of LexA(ind-) allele expression on growth (C) Expression of a $P_{dinC}$-YFP reporter as a function of time after addition of MMC in a genetic background including the IPTG-inducible $P_{hs}$-lexA(ind-) allele. Shown are results with 0 IPTG (no induction, orange) and 100μM (full induction, purple). Circles represent individual measurements, lines represent best linear fits and shaded area the boundary of error.

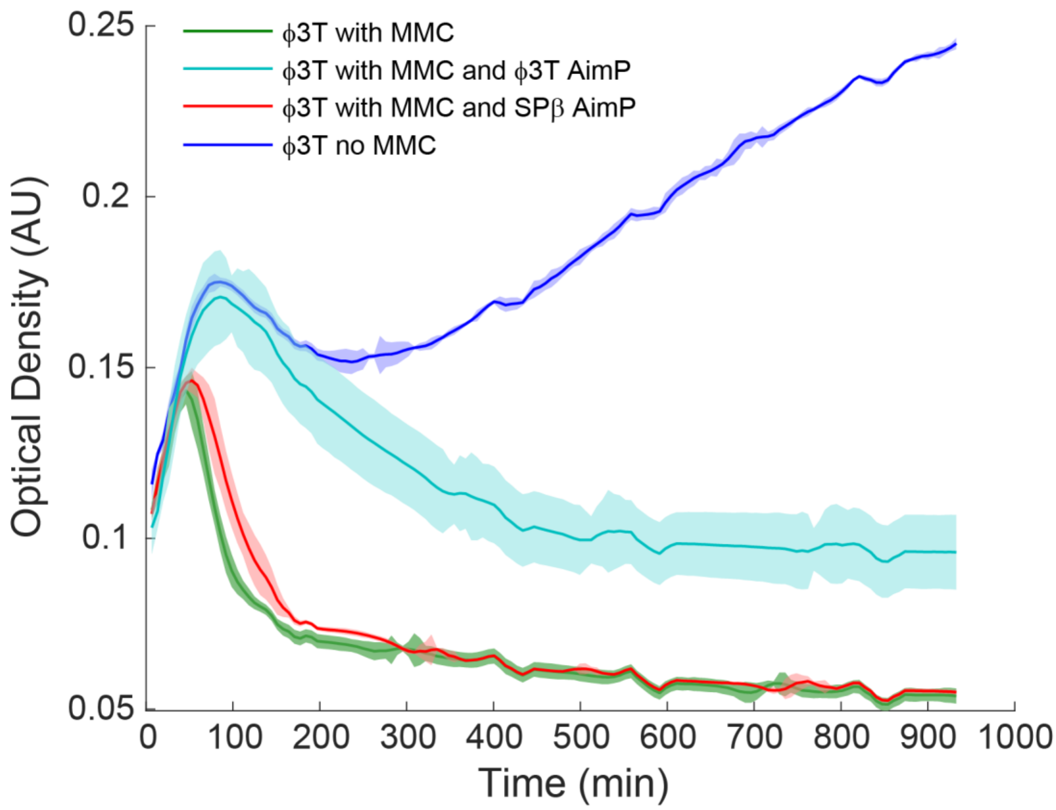

**Supplementary figure 9: ϕ3T induction is repressed by its AimP**. Shown are graphs of optical density as a function of time for four cultures of a ϕ3T lysogens grown in LB broth, as described in the legend. Solid lines mark the mean of three replicates done on the same multi-well plate. The region around each line indicates the standard error of the mean. ϕ3T AimP is the ϕ3T arbitrium peptide (SAIRGA), and SPβ AimP is the peptide GMPRGA.

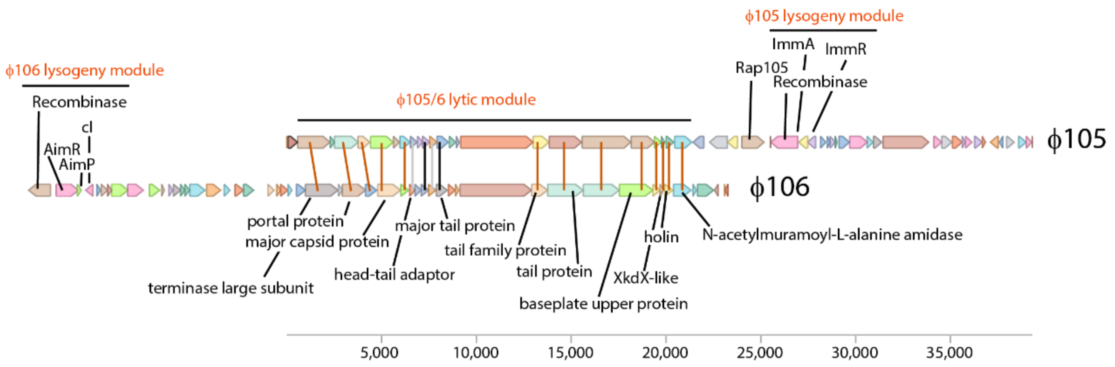

**Supplementary figure 10: Comparison of phages $\phi$105 and $\phi$106.** We use the following sequences for comparison. $\phi$105 is analyzed using its directed sequence file at the NCBI accession number NC_048631. $\phi$106 is defined here as the lysogenic segment of *Bacillus subtilis subsp. inaquosorum strain KCTC 13429* (NCBI accession number NZ_CP029465) from locus tag DKG76_RS14125 to locus tag DKG76_RS14370. $\phi$106 proteins were blasted against $\phi$105 proteins. Marked are proteins with high (red, Expectation<$10^{-20}$), medium (black, $10^{-10}$<Expectation<$10^{-20}$) and few low homology genes within the lytic module (gray, $10^{-5}$<Expectation<$10^{-10}$). There is little or no homology out of the region annotated here as the lytic module. Protein annotation within the lytic module is based on phage $\phi$105 annotation. The lysogenic module of phage $\phi$105 including the recombinase $ImmRA_{105}$ and Rap gene are shown [6]. Homologs of these genes are absent from $\phi$106. Also shown is the putative lysogenic module of $\phi$106 analyzed in this work, which is absent from phage $\phi$105.

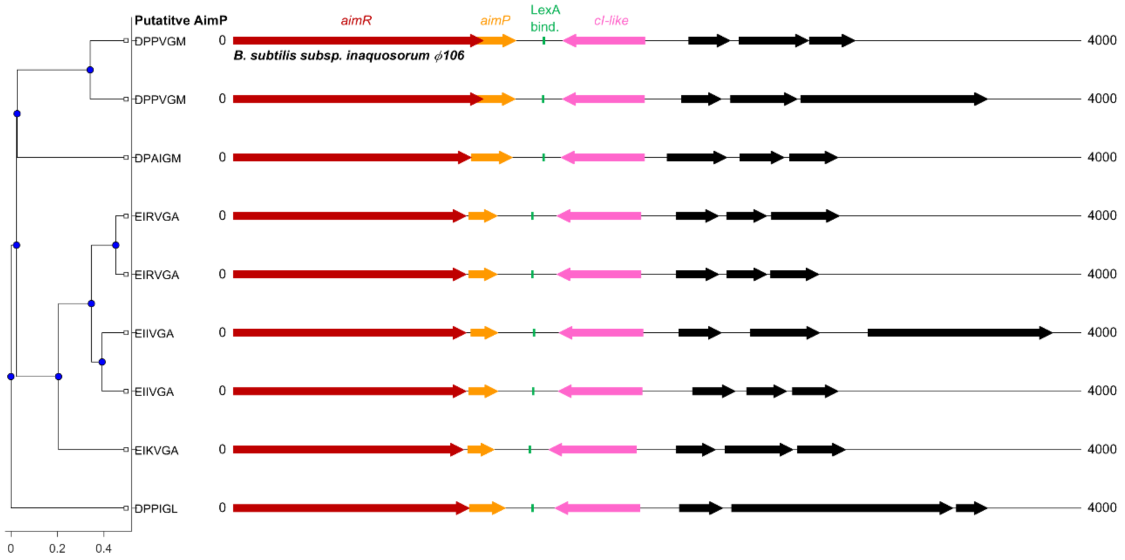

**Supplementary figure 11: Consistency and diversity of the ϕ106 family putative lysogeny module.** Left: Shown is the phylogenetic tree of 8 different AimR clade 1 proteins (out-grouped by AimR from phage SPβ). Each of the eight corresponding *aimR* genes has an adjacent *aimP* gene coding for the putative mature arbitrium AimP signal shown to the right of the corresponding tree leaf. Right: the genomic organization of the arbitrium locus is shown for each AimR gene. Colored arrows represent genes, according to the annotation at the top. Black arrows correspond to additional genes. Green line marks the position of the putative LexA binding site. The ϕ106 system used in this work is specifically marked (strain *inaquosorum*). The NCBI accession number of the AimR proteins (and the genome GCF number) as shown in the phylogeny, from top to bottom, are:

WP_003237457 (003148415), WP_101605414 (002850535), ARW33040 (001747445), WP_049627412 (004119735), WP_047936208 (001023595), WP_003220312 (005218185), API45091 (002982175), WP_017417251 (003860445), WP_181217268 (013620725).

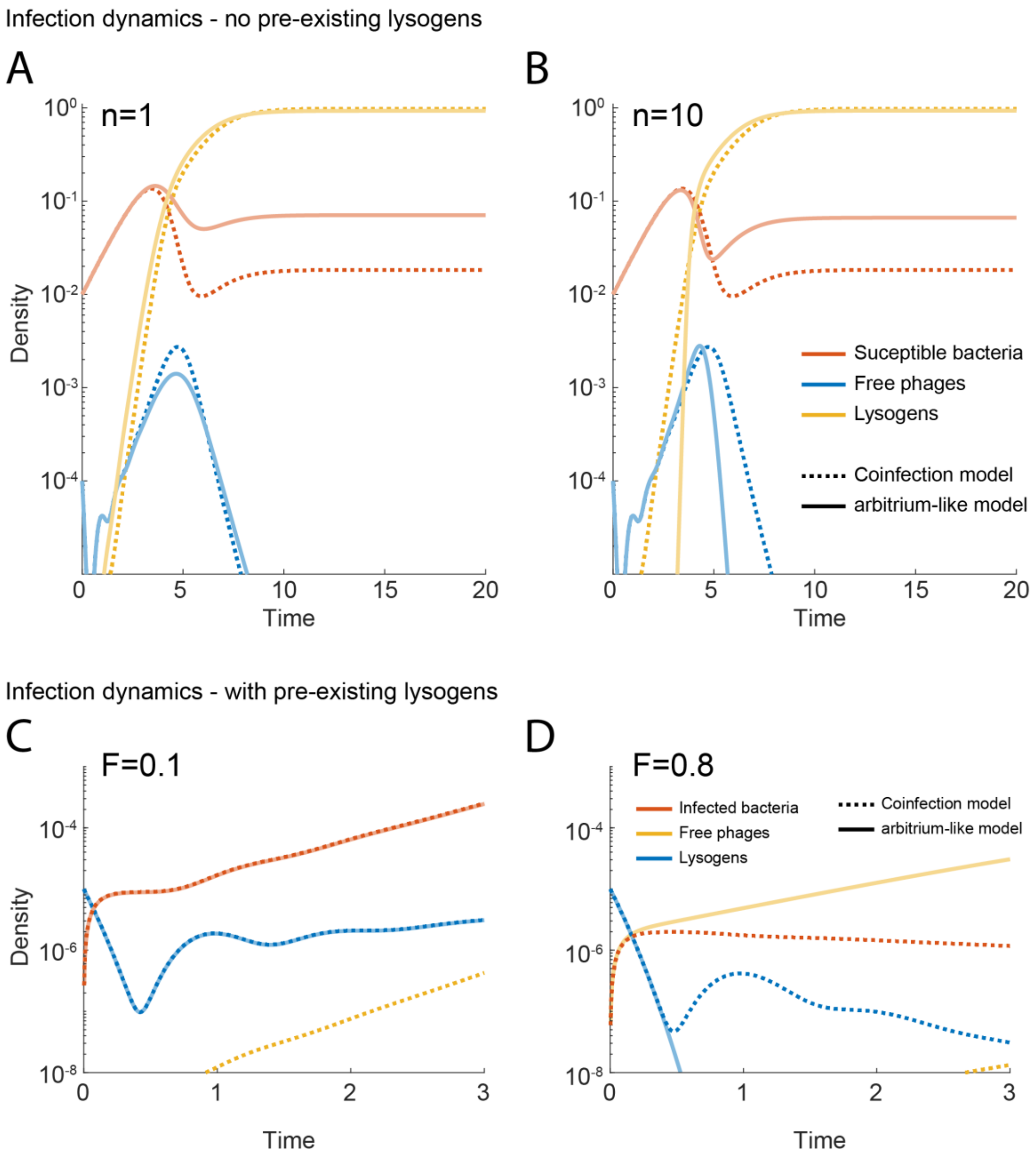

**Supplementary figure 12: Infection dynamics for the two phage models with and without an initial Lysogenic population.** (A,B) The two models show similar infection dynamics in the absence of an initial lysogen population. Shown are solutions of eqs. 5-10 (coinfection model) and 11-16 (arbitrium-like model) of Supplementary text. In both panels the parameters are: $\gamma = 1, \eta = 1000, \omega = 1.5, \beta = 4, \kappa = 1, r = \frac{1}{3}\omega^{-1}, c = 0.01, \sigma = 300, \lambda = 300, S_{th} = 0.4$. Initial phage and permissive bacteria levels are $P(0) = 10^{-4}, B(0) = 10^{-2}$. The difference between the panels is the hill coefficient of the response to signal in the arbitrium-like model which is (A) $n = 1$ and (B) $n = 10$. Shown are values for free phages (blue), permissive bacteria (red) and lysogens (orange) for the two models – coinfection sensing (dashed lines) and arbitrium-like (solid lines). (C,D) The two models differ in infection dynamics in the presence of lysogens. Simulations of the same equations with the same parameters as in A,B, except for the following differences, $P(0) = 10^{-5}, B^{tot}(0) = 0.01,\ n = \infty$ (threshold response), $S_{th} = 0.28$ (optimal). The panels differ in the initial fraction of pre-existing Lysogens (A) $F = 0.1 - B(0) = 0.9B^{tot}(0); L_o(0) = 0.1B^{tot}(0)$. (B) $F = 0.8 - B(0) = 0.2B^{tot}(0); L_o(0) = 0.8B^{tot}(0)$.

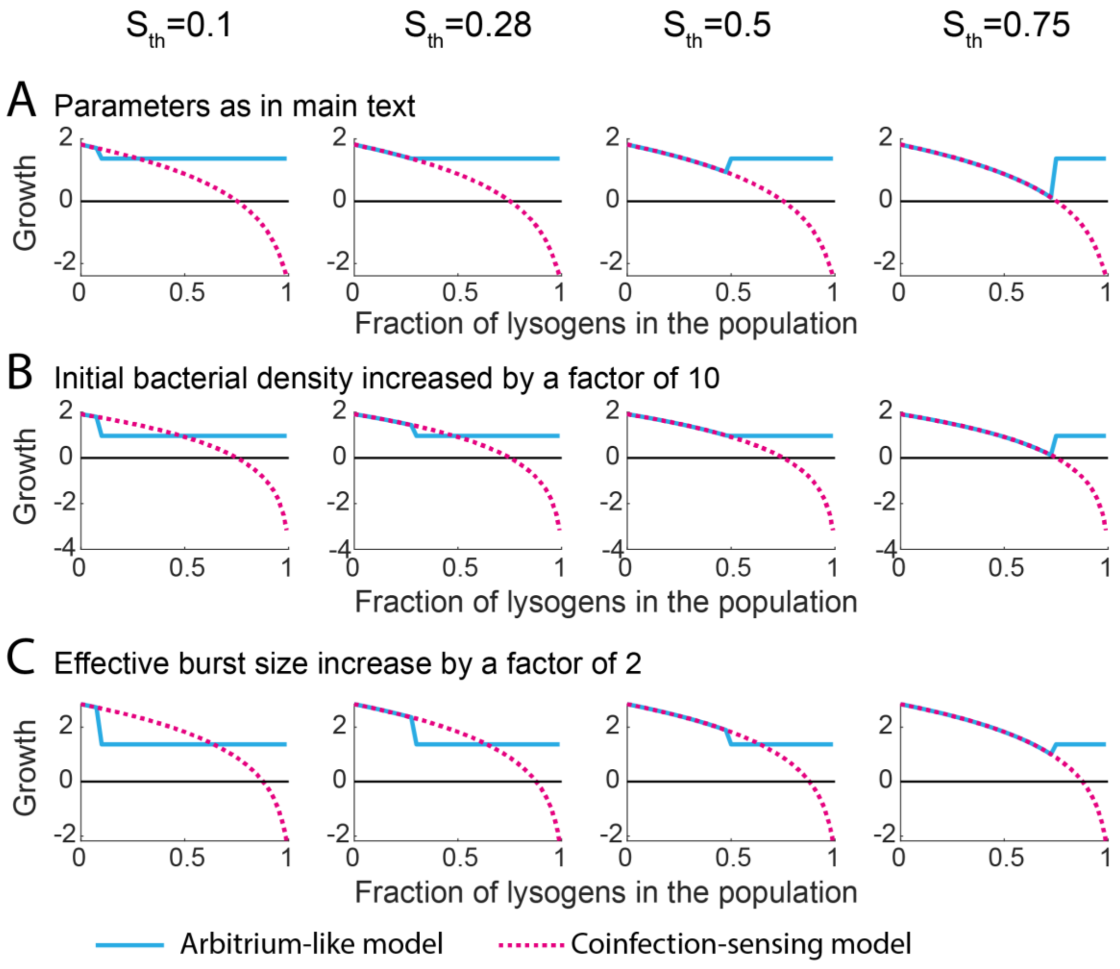

**Supplementary figure 13: Phage growth rates of the two models for different parameters.** Shown are the average growth rates of phages as a function of frequency of lysogens for the coinfection-sensing (dashed line, eqs. 5-10 in Supplementary text) and arbitrium-like (solid line, eqs. 11-16 in Supplementary text) models for different parameters. Phage average growth rate is defined in the Supplementary text (eq. 18). (A) The values used for the simulations are $\gamma = 1, \eta = 1000, \omega = 1.5, \beta = 4, \kappa = 1, r = \frac{1}{3}\omega^{-1}, c = 0.01, \sigma = 300, \lambda = 300, n = \infty$. Initial phage levels are $P(0) = 10^{-5}, B(0) = 10^{-2}$. The columns represent different signal threshold frequencies $S_{th}$ as indicated. (B) as in (A), but with $B(0) = 10^{-1}$. (C) As in (A), but with $\beta = 8$.

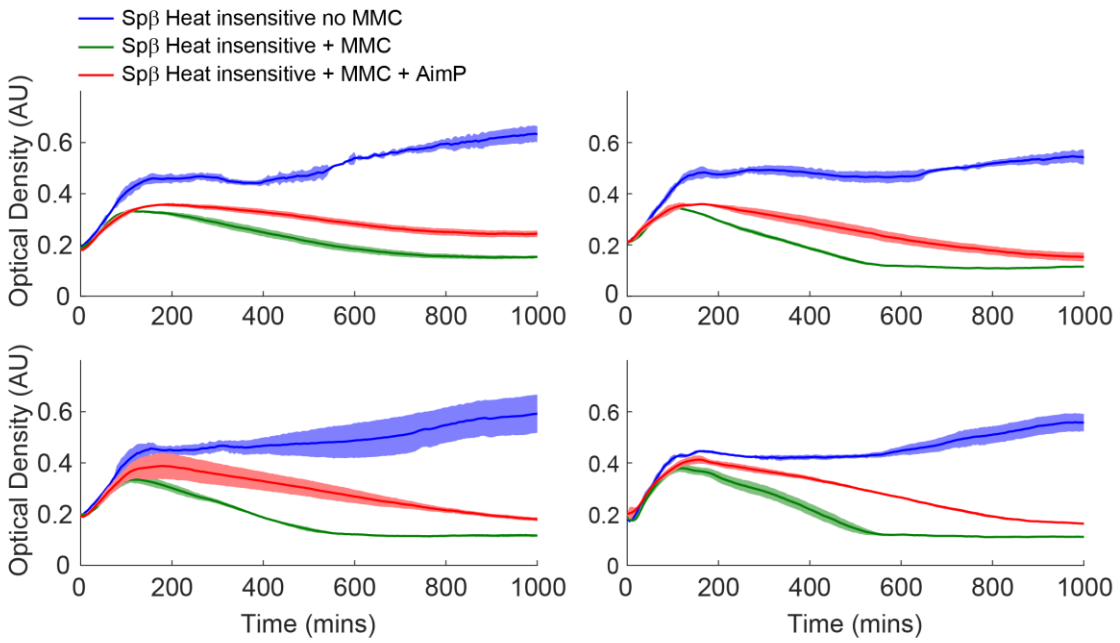

**Supplementary figure 14: The heat insensitive SPβ variant responds to MMC and AimP at 37$^0$.** Shown are four repeats of plate reader readings of optical density as a function of time of the various strains and conditions described in the legend. Cells were grown in LB. The heat insensitive variants of SPβ lyse upon addition of MMC and this lysis is reduced upon addition of AimP, similarly to what is found in the heat-sensitive strain (Supplementary Fig. 1).

### Supplementary text: The impact of prophage signaling on infection dynamics: mathematical analysis

The aim of this analysis is to compare the impact of lysogens on infection dynamics in a scenario where phage density is measured through co-infection (as in phage $\lambda$), and in a scenario where both lysogens and infecting phages can signal their presence, as in the arbitrium-coding phages we have characterized in this work.

We wish to demonstrate three major points:

1. In the absence of prior existing lysogens, infection dynamics are similar in the two scenarios.
2. In the presence of lysogens, signaling provide a fitness benefit to phages, especially if it is tuned to an optimal frequency of lysogens in the population. This adaptive benefit is true both for control of lysis-lysogeny and control of phage induction.
3. In structured environments, the benefits of lysogenic signaling are even larger.

To demonstrate these points, we first present a mathematical model of phages using either coinfection sensing or arbitrium-like signals to control lysis-lysogeny (and induction). We then use these models to address all three points.

#### Modeling infection dynamics for the two sensing mechanisms

We consider two models of temperate phage infection. The first assumes that lysogenization rate upon infection is regulated by the number of phages co-infecting a cell within a certain time window (Co-infection mechanism). This model generally captures the biology of $\lambda$-phage infection and is similar to other models used to describe it[7–9]. The second assumes that lysogenization probability is dependent on the level of signal sensed by an infecting phage, as described for the arbitrium system, but including the observation made in this work that lysogens also produce the signal (arbitrium-like mechanism)[10]. It is worth noting that the two mechanisms can co-exist, but for the sake of simplicity we would assume that phages with arbitrium-type sensing do not have an additional coinfection-based regulation.

We further assume that except for these differences, the two types of phage have the same infection dynamics parameters, that is their infectivity, eclipse time and burst size are identical. The equations describing infection dynamics would therefore be the same, but for lysogenization. The variables used in the model are those for the densities of permissive bacteria ($B$), phages ($P$), infected cells ($W$) and lysogens ($L$). In the signaling model, we also keep record of extracellular signal concentrations ($S$). We note that in both models, we divide the infected cell population into $N$ compartments with serial transfer between compartments. This is equivalent to adding the eclipse time as a time-delay.

Below we write down and explain the two models. The two models are based on the same set of equations for the lytic part, and we therefore write it down first with appropriate explanations. We then write down the equations for the two mechanisms. Common parts of the equations are marked in black and model specific parts are colored.

All equations are solved numerically using Matlab

#### Lytic model

The equations describing the lytic core of the two model are the following:

| Parameters used | |
|---|---|
| $\eta$ | Infection rate |
| $\gamma$ | Growth rate |
| $\omega$ | Eclipse rate |
| $\beta$ | Burst size |
| $\kappa$ | Carrying capacity |
| $r$ | Lysogeny time-window |
| $c$ | Cost of lysogeny |
| $\sigma$ | Signal production |
| $\lambda$ | Signal uptake |
| $S_{th}$ | Threshold signal response |
| $n$ | Signal response sensitivity |

1. $\frac{dB}{dt} = \gamma \cdot B \cdot \left(1 - \frac{B_{tot}}{\kappa}\right) - \eta \cdot P \cdot B$ – permissive bacteria levels are regulated by bacterial growth and phage infection. We assume that growth is logistic (the $\left(1 - \frac{B_{tot}}{\kappa}\right)$ term), where $B_{tot} = B + L + \sum W_i$.
2. $\frac{dW_1}{dt} = \eta \cdot P \cdot B - N\omega \cdot W_1$ – Infected bacteria are the product of infection, infection moves to the next compartment with a rate which is $N$ times faster than its eclipse rate. Since there are $N$ compartments, this provides a total eclipse rate $\omega$ for the whole process, but would effectively introduce this as a time-delay. We assume here that the infected, non-lysogenic, cells do not grow.
3. $\frac{dW_k}{dt} = N\omega(W_{k-1} - W_k); k = 2, \dots, N$ – Transmission through infection compartments.
4. $\frac{dP}{dt} = \beta\omega \cdot W_N - \eta \cdot P \cdot B_{tot}$ – Free phages form with a given burst size $\beta$ from the late infected cells. Free phages are consumed by infecting any type of cell.

#### Lysogenization through co-infection

In the co-infection lysogenization model we assume that there is a time window $\tau = \frac{r}{\omega}$ where the infection of an infected cells by a second phage would lead to lysogenization. This is a simplified assumption, as in principle co-infection leads to changes in the probability of infection. In such a case, the equations would be:

5. $\frac{dB}{dt} = \gamma \cdot B \cdot \left(1 - \frac{B_{tot}}{\kappa}\right) - \eta \cdot P \cdot B$
6. $\frac{dW_1}{dt} = \eta \cdot P \cdot B - N\omega \cdot W_1 - \eta \cdot P \cdot W_1$
7. $\frac{dW_k}{dt} = N\omega(W_{k-1} - W_k) - \eta \cdot P \cdot W_k; k = 2, \dots, j_\tau; \frac{j_\tau}{N} = r$
8. $\frac{dW_k}{dt} = N\omega(W_{k-1} - W_k); k = j_{\tau+1}, \dots, N$
9. $\frac{dP}{dt} = \beta\omega \cdot W_N - \eta \cdot P \cdot B_{tot}$
10. $\frac{dL}{dt} = \gamma \cdot (1 - c) \cdot L \cdot \left(1 - \frac{B_{tot}}{\kappa}\right) + \eta \cdot P \cdot \sum_1^{j_\tau} W_k$

#### Lysogenization through Signaling (arbitrium-like mechanism)

Here, we assume that infected cells and lysogens produce a signal at a certain production rate. We assume that the signal is taken up by the cells at a constant rate, as has been shown for RRNPP-type quorum-sensing system [11,12]. Though the response is dependent on the intracellular level of the signal, we will assume for simplicity that it depends on the extracellular concentration of the signal. Lysogenization probability is assumed to be a hill function of the signal level and to occur immediately upon infection. The equations describing this process are:

11. $\frac{dB}{dt} = \gamma \cdot B \cdot \left(1 - \frac{B_{tot}}{\kappa}\right) - \eta \cdot P \cdot B$

12. $\frac{dW_1}{dt} = \left(1 - f(S)\right) \cdot \eta \cdot P \cdot B - N\omega \cdot W_1; f(S) = \frac{S^n}{S^n + S_{th}^n}$

13. $\frac{dW_k}{dt} = N\omega(W_{k-1} - W_k); k = 2, \dots, N$

14. $\frac{dP}{dt} = \beta\omega \cdot W_N - \eta \cdot P \cdot B_{tot}$

15. $\frac{dL}{dt} = \gamma \cdot (1 - c) \cdot L \cdot \left(1 - \frac{B_{tot}}{\kappa}\right) + f(S) \cdot \eta \cdot P \cdot B$

16. $\frac{dS}{dt} = \sigma(L + W) - \lambda \cdot B_{tot} \cdot S$

Signal concentration in RRNPP type quorum-sensing systems is dependent on production and uptake of the signal, both dependent on cell density. This results in a concentration which is more clearly dependent on the frequency of producers in the population, than on the density of cells (at least above a certain density) [11]:

17. $S_{st} \sim \frac{\sigma}{\lambda} \frac{(L+W)}{B_{tot}}$

We therefore set the threshold signal concentration to be a fraction of the maximal allowed concentration, $S_{th} = \theta \frac{\sigma}{\lambda}$. The response function is simplistic, as it assumes no "leakiness" of lysogenization. This assumption is also made in the co-infection model.

#### Normalization, relevant time scales and parameters choice

The equations can be normalized by setting bacterial density so that $\kappa = 1$, by setting time so that $\gamma = 1$ and by setting the signal production rate per cell to equal the signal uptake rate per cell, making the maximal $S_{st}$ to be 1.

Different processes occur at different time-scales and the nature of the infection dynamics would depend on the relation between these time scales. We list the relevant time-scales and discuss the values we have defined for each parameter:

- Bacterial growth is characterized by the time scale $\gamma^{-1}$. In normalized units $\gamma = 1$.
- Eclipse time is characterized by $\omega^{-1}$. Phage initial growth rate would equal $\frac{\omega^{-1}}{\log(\beta)}$. We assume that the eclipse time is of the same order of the cellular growth rate. In normalized units we choose $\omega = 1.5$.
- Burst size. The real burst size of phages is of the order of tens to few hundreds. However, we choose a significantly lower number representing an "effective" burst size, taking into account loss of phages in the environment (degradation, infection of "wrong" bacteria etc). This may vary a lot and we will explore values between $\beta \sim 1.5 - 10$.
- Infection time is the time it takes a free phage to infect a bacterium. This is equal to $(\eta B)^{-1}$. In normalized units, the maximal bacterial density is 1, leading to an infection time of $\eta^{-1}$. We assume that this is much lower than the growth rate. We choose $\eta = 1000$, so that even for $B = 0.01$ as an initial condition, this would be significantly lower than the eclipse time.
- Signal equilibration time is equal to $(\lambda \cdot B_{tot})^{-1}$. In normalized units the maximal level of $B_{tot}$ is 1, and the maximal signal equilibration time-scale is $\lambda^{-1}$. We assume that this is significantly shorter than the growth rate even for intial infection dynamics. We therefore choose $\lambda = 300$. To normalize the level of signal, we assume that $\sigma = \lambda$, which implies that the maximal level of signal possible is $S_{st} = 1$. For the response we use either a linear response ($n = 1$) or a threshold response ($n = 10$) and varying threshold. Note that the level of signal is proportional to a large extent to the frequency of lysogens and infected cells.
- Lysogeny waiting time for the coinfection model is assumed to be $\frac{1}{6}$ of the eclipse time. Cost of lysogeny is assumed to be 0 for simplicity.

#### Infection dynamics for coinfection and arbitrium-like sensing mechanisms are similar.

Numerical results for the above-mentioned set of parameters are shown in Fig. S12A,B for two cases – a linear response of lysogeny to the signal ($n = 1$) and a threshold response ($n = 10$). The bacteria and phage initial conditions are set to $B(0) = 0.01$; $P(0) = 10^{-4}$. The two cases are compared to a coinfection model with the same values. One can see that the infection dynamics are very similar for the linear signal response case. In this case, the chances of a second phage entering into an infected cell, which corresponds to a switch to lysogeny, is correlated with the phage density which is proportional to signal density. In the case of a threshold response, the formation of lysogens in the signaling model occurs naturally later and more abruptly than in the coinfection model.

#### Early infection in the presence of lysogens

To model infection dynamics of phages in the face of an existing lysogen population, we do the following modification to the above equations (for both mechanisms). We mark the existing lysogen population with a different variable $L_o$. We assume that in both mechanisms, this population is growing with the same growth rate as other lysogens, and that it is not permissive to effective infections, thus serving as a sink for infecting phages. For the arbitrium-like mechanism, we assume that these lysogens also secrete and uptake signal at the same rates as infected cells. We assume a very low initial phage density $P(0) = 10^{-5}$, leading to a very low probability for coinfections.

As a fraction $F$ of the population is lysogenic, we expect that only a fraction $(1 - F)$ of the initial phage population would infect permissive host cells and lead to further lysis or lysogeny. We therefore defined the average growth rate until time $t$ of the phages to be:

18. $G(t) = \frac{1}{t}\log\left(\frac{P^T(t)}{(1-F)P(0)}\right)$, where $P^T = P + L + \sum_1^N W_i$

The data shown in Fig. 3D of the manuscript shows this average growth rate at time $t = 4$ as a function of the initial lysogen fraction $F$. In Supplementary Fig. 12C,D we show the trajectory of free phages, infected cells and new lysogens (excluding the initial $L_o$ population) as a function of time for infection at $F = 0.1$ and $F = 0.8$ for the coinfection sensing and arbitrium-like mechanisms. As can be seen, phages with the coinfection sensing mechanism are essentially fully lytic, as there are no coinfections. The arbitrium-like mechanism is also fully lytic at $F = 0.1$, but is fully lysogenic at $F = 0.8$, where lysogeny is a superior strategy to lysis.

In Supplementary Fig. 13, we explore the dependence of $G$ on the initial bacterial density, the effective burst size and the threshold frequency. In general, phages with arbitrium-like mechanism behave like those with a coinfection sensing mechanism below the signaling threshold where both are lytic. Their relative growth is always better above the signaling threshold and herd immunity, as they have positive growth while the lytic strategy of the coinfection-sensing phages leads to decline. Under some conditions, there is a range where coinfection-sensing has better growth, if the signaling phages switch to lysogenization at a fraction of lysogens for which lytic growth is still superior.

This argument regarding infective phages can also be extended to the regulation of induction, as it is not worth-while to induce if the resulting lytic growth of the outcoming phages is less than the possible benefit of lysogenic growth. We therefore expect that regulation of induction would depend similarly on lysogen frequency in signaling-based phages. The dependence on DNA damage has been explained elsewhere and is still valid for this case as well [8].

Finally, the transition graph could be understood analytically with a simplified model which assumes no growth of the bacteria. In this case, the lytic growth per phage life cycle is simply $G(F) = R_0\omega^{-1}(1-F)$, where $R_0$ is the basic reproductive number and $\omega^{-1}$ is the eclipse time. If the lysogenic growth is $G_L < R_0\omega^{-1}$, the optimal strategy is to be lytic until $F_{th} = 1 - \frac{G_L}{R_0}$ and then switch to lysogeny.

#### Adaptivity of signaling in structured populations

A structured population is characterized by an uneven (correlated) distribution of genotypes [13]. The local nature of infections in colonies and biofilms and the micron range of phage signaling in these communities [12], imply the fraction of lysogens in a given signaling range is expected to be either very high ($F \sim 1$) – a microcolony of lysogens, or very low $F \ll 1$ – a microcolony of susceptible bacteria. Furthermore, it was shown in dense populations, phage signaling acts on a micron scale[12]. Under these assumptions, the benefit of lysogenic signaling are sharpened:

1. For $F \ll 1$ both arbitrium-like and coinfection sensing mechanisms would yield an initial lytic expansion, which will be the optimal strategy.
2. For $F \sim 1$, it is most likely that the community is beyond the herd immunity frequency and therefore the lysogenic strategy adopted by the arbitrium-like mechanism at high frequencies is better than the lytic strategy of the coinfection sensing mechanism.

Structured populations provide a lower probability for mixed populations with an intermediate frequency of lysogens, where either mechanism can be more adaptive, depending on infection parameters. Structured populations thereby increase the adaptive value of the arbitrium-like mechanism which is always more adaptive at the extremes of lysogen frequencies.
